# Tissue-specific changes in the RNA structurome mediate salinity response in *Arabidopsis*

**DOI:** 10.1101/604199

**Authors:** David C. Tack, Zhao Su, Yunqing Yu, Philip C. Bevilacqua, Sarah M. Assmann

## Abstract

RNA structures are influenced by their physico-chemical environment. Few studies have assessed genome-wide impacts of abiotic stresses on *in vivo* RNA structure, however, and none have investigated tissue-specificity. We applied our Structure-seq method to assess *in vivo* mRNA secondary structure in Arabidopsis shoots and roots under control and salt stress conditions. Structure-seq utilizes dimethyl sulfate (DMS) for *in vivo* transcriptome-wide covalent modification of accessible As and Cs, i.e. those lacking base pairing and protection. Tissue type was a strong determinant of DMS reactivity, indicating tissue-specificity of RNA structuromes. Both tissues exhibited a significant inverse correlation between salt stress-induced changes in transcript reactivity and changes in transcript abundance, implicating changes in RNA structure and accessibility in transcriptome regulation. In mRNAs wherein the 5’UTR, CDS and 3’UTR concertedly increased or decreased in mean reactivity under salinity, this inverse correlation was more pronounced, suggesting that concordant structural changes across the mRNA have the greatest impact on abundance. Transcripts with the greatest and least salt stress-induced changes in DMS reactivity were enriched in genes encoding stress-related functions and included housekeeping functions, respectively. We conclude that secondary structure regulates mRNA abundance, thereby contributing to tissue specificity of the transcriptome and its dynamic adjustment under stress.

**One Sentence Summary:** Transcriptome-wide methods reveal dynamic tissue-specific and salt stress-dependent modulation of mRNA accessibility and structure, and correlated mRNA abundance changes.

## Introduction

Given a predicted world population of 9.1 billion by 2050, improvements in crop stress tolerance will be critical to meet required increases in food production (FAO, 2009). Salinity stress is one of the most common environmental stresses affecting crop productivity worldwide (Munns and Tester, 2008). Elevated soil salinity affects 932 million hectares (Rengasamy, 2006) of agricultural fields (roughly the size of the entire United States) and causes US$ 27.3B in crop losses annually (Qadir et al., 2014). Soil salinization arises from irrigation practices and, increasingly, from seawater inundation as a result of climate change, which, e.g. threatens lowland rice production in many regions of Asia (Vu et al., 2018). In this context, an urgent goal is to better understand plant responses to salinity stress, especially at the molecular level. Saline soils cause cellular dehydration by thermodynamically opposing plant water uptake, and Na^+^ and Cl^−^ accumulation in plant tissues negatively affects plant growth by hampering metabolic processes, promoting oxidative stress (Abogadallah, 2010), and reducing photosynthetic efficiency (Tester and Davenport, 2003; Deinlein et al., 2014). Cells are tiny but accurately controlled and complex machines; as cytosolic solute composition changes, cellular molecules, including DNA, RNA, proteins, and metabolites, may function differently from their normal status due to changes in folding and interaction. In particular, salt generally enhances RNA folding and disrupts RNA-protein interactions (Bloomfield et al., 2000).

Several studies have reported dramatic impacts of salt stress on the Arabidopsis transcriptome (Kreps et al., 2002, Anderson et al., 2018). Salinity also leads to marked transcriptome changes in other plant species, including important crops such as rice (Wang et al., 2018), tomato (Sun et al., 2010), barley (Bahieldin et al., 2015), sorghum (Cui et al., 2018), and cotton (Zhu et al., 2018), demonstrating the universal impact of salinity and the importance of salt-induced effects on mRNA abundance. Transcript abundance reflects the balance between mRNA transcription and mRNA turnover, and post-transcriptional regulation is known to play vital roles in plant responses to environmental factors (Floris et al., 2009). For example, salt stress changes alternative splicing patterns (Feng et al., 2015) and mRNA decay pathways as assessed in Arabidopsis and other species (Kawa and Testerink, 2017), which provides layers of regulation on the transcriptome and encoded proteome (Floris et al., 2009).

While transcriptome analyses often center on measuring changes in abundance, changes in mRNA secondary and tertiary structures influence many aspects of post-transcriptional regulation, including alternative polyadenylation, splicing, stability, localization, degradation and translation (Bevilacqua et al., 2016). RNA structure *in vitro* is well-known to be affected by metal ions and by proline (Lambert and Draper, 2007), a compatible solute synthesized during salt stress, but to what extent salt stress affects mRNA structure *in vivo* and genome-wide, with attendant implications for post-transcriptional regulation, has not been studied in any organism.

Numerous chemicals have been identified that react with distinct regions of the RNA bases and sugar-phosphate backbone (Mitchell et al., 2019; Bevilacqua & Assmann 2018; Leamy et al., 2016; Bevilacqua et al., 2016; Wilkinson et al., 2006; Ziehler and Engelke, 2000). These chemicals covalently modify RNA at the Watson-Crick (WC) face, the non-WC face or the ribose hydroxyl group. We developed Structure-seq (Ding et al. 2014) as a transcriptome-wide method that currently utilizes dimethyl sulfate (DMS) (Tijerina et al., 2007) as a chemical probe of RNA structure *in vivo* and genome-wide. DMS methylates unpaired and unprotected adenosine (A) and cytosine (C) at the imino nitrogen on the WC face, which blocks subsequent reverse transcription. The resulting truncated complementary DNAs (cDNAs) are ligated to adaptors and analyzed by next-gen sequencing methods. The readout of this method is a reverse transcription stop one position prior to each DMS-modified nucleobase. We recently improved Structure-seq experimentally in Structure-seq2 (Ritchey et al., 2017) and computationally in StructureFold 2 (Tack et al., 2018).

In a recent application of Structure-seq, we uncovered a significant negative correlation between changes in DMS reactivity and changes in mRNA abundance following brief heat shock in rice, suggesting that mRNA unfolding promotes access to degradative processes (Su et al. 2018). Here, we have used Structure-Seq2 (Ritchey et al. 2017) coupled with the StructureFold2 computational pipeline (Tack et al., 2018) to resolve *in vivo* structuromes of Arabidopsis shoot and root tissue under both unstressed and salt stressed conditions. Our results reveal tissue-specific and stress-modulated aspects of the RNA structurome.

## Results

### Experimental Design and NaCl Treatment

To investigate the effect of salt stress on the RNA structurome, we applied our Structure-seq2 protocol (Ritchey et al., 2017; Su et al., 2018) to 24-day-old hydroponically grown Arabidopsis with and without treatment with 100 mM NaCl for 48 h. We confirmed the effectiveness of our salt stress by observation of reduced rosette growth (Figure 1A vs. B) and by significant increases in proline content in our salt-stressed plants (Figure 1C); proline is commonly accumulated as a compatible solute following salt stress (Trinchant et al., 2004; Mattioli et al., 2009). We also detected significantly increased Na^+^ content in both shoots and roots of NaCl-treated plants (Figure 1D). Root tissue often shows decreased K^+^ content post-salt stress (Sun et al., 2015), a phenomenon we also observed (Figure 1E) (Yu and Assmann, 2015). Together, these observations confirm the efficacy of our NaCl treatments. All accompanying measurements and statistics, including those of additional ions, can be found in Table S.1 and analysis of this via ANOVA can be found in Table S.2.

**Figure 1.**
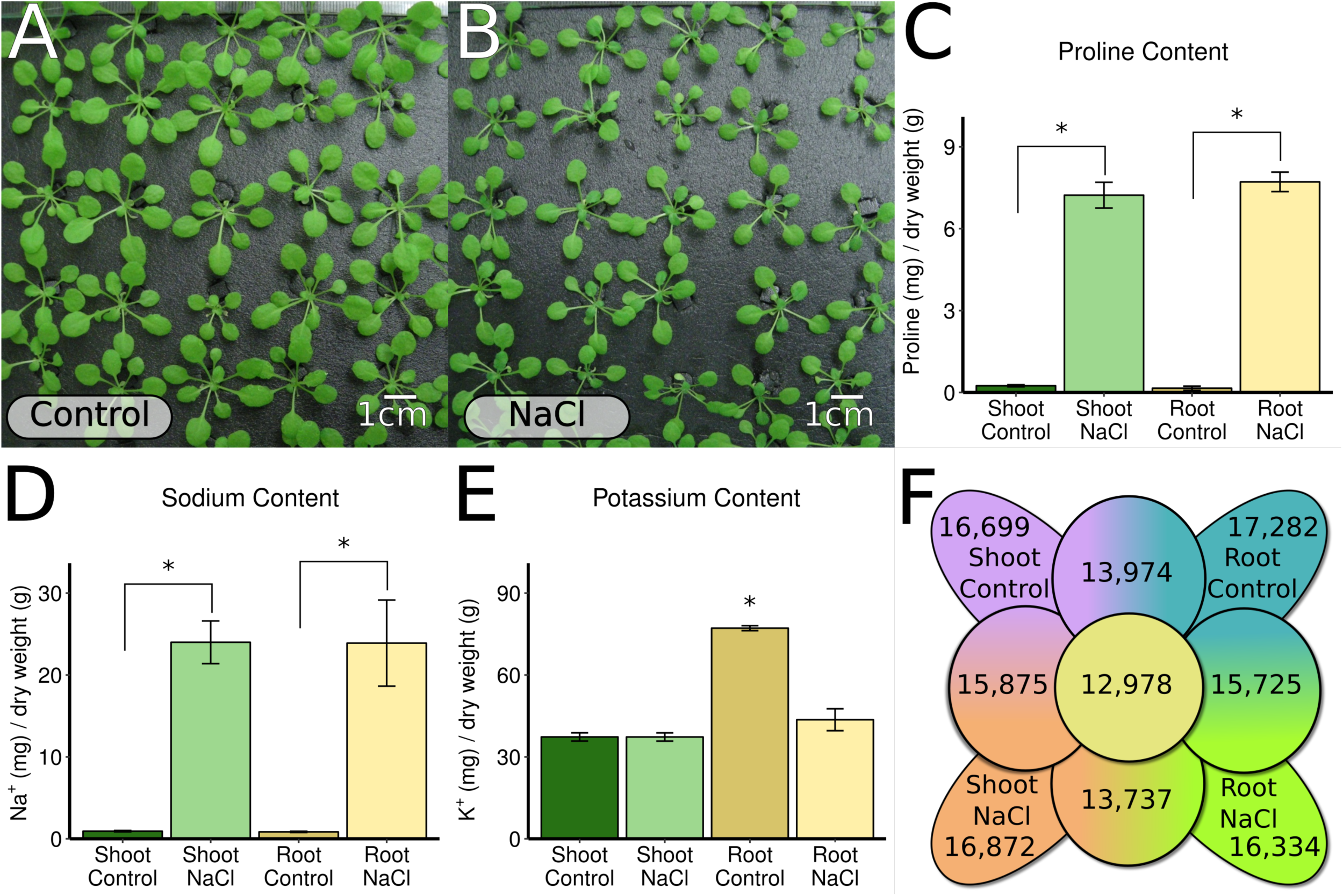
The Effect of Salt Stress on Arabidopsis Phenotypes and Ion Content. A,B) 24 day old hydroponically grown Arabidopsis (A) under control conditions and (B) after exposure to 48h of 100 mM NaCl stress. C, D, E) proline, sodium, and potassium content of both control and NaCl stressed plants; NaCl stress greatly increases the proline and sodium content of both shoots and roots, while lowering the potassium content of roots. All accompanying measurements and statistics, including those for additional ions, can be found in Table S.l and analysis via ANOVA can be found in Table S.2. * indicates p < 0.05. F) Total number of transcripts with Structure-seq coverage within and between each tissue and condition surveyed.

### Structure-seq Library Preparation and Evaluation

Custom Structure-seq libraries (three independent biological replicates per condition) were generated from control and salt stress conditions. Libraries were prepared from poly(A) RNA isolated separately from shoot and root tissues after treatment with DMS or after a -DMS control treatment, yielding a total of 24 libraries (Table W.1). We mapped our reads to a set of TAIR 10 cDNAs (See Methods, rRNAs removed), yielding an average of 90.3% of reads mapping to the transcriptome in all libraries (Table W.2). Reads that failed to meet mapping criteria (see Methods) or mapped to the incorrect strand were discarded (Table W.3). Reverse transcription (RT) stops in the +DMS libraries were largely specific to adenines (As) and cytosines (Cs) (Table W.4), as expected. The DMS-induced RT stops in the replicate libraries under each condition were highly reproducible (mean r=0.95) as shown by high correlations of the number of RT stops on each nucleotide between any two +DMS libraries (Figure S.2). Given the high reproducibility, library data from the three replicate libraries were merged as the final datasets for downstream analyses (Ding et al., 2014). We define coverage as the number of sequenced RT stops per each immediately upstream A and C present in each transcript in the pool of all +DMS biological replicates of a condition; thus, a coverage threshold of ≥ 1 mandates that we sequenced an average of one or more RT stops for each A or C residue of a given transcript within a condition in order to consider the structural information for that transcript to be resolvable (Kertesz et al., 2010).

Ultimately, we had 16,699 and 16,872 mRNAs that met our criteria to derive reliable DMS reactivity and mRNA secondary structures under control and salt conditions in shoot tissue, and 17,282 and 16,334 corresponding mRNAs in root tissue (Figure 1F). Among these, 15,875 shoot mRNAs and 15,725 root mRNAs had sufficient coverage under both control and salt stress conditions, which allowed assessment of salinity effects on mRNA structure in each tissue (Figure 1F). There were 13,974 mRNAs that had sufficient structural coverage in both root and shoot in control conditions and 13,737 mRNAs that had sufficient structural coverage in both root and shoot under salt stress, which allowed us to investigate the tissue specificity of mRNA structure, and structural responses to salinity in each tissue. Among all four data sets, a remarkable 12,978 mRNAs had sufficient coverage in all tissues and conditions, allowing for structural comparison of a set of common mRNAs

### Salt Stress Alters the mRNA Structurome

We compared the distributions of mean reactivity (Figure 2, Tables 1, S.3.a) of whole transcript data (Figure 2A, 2E) as box-violins in both control (left violin, blue) and NaCl (right violin, yellow) conditions while indicating a change in mean reactivity between conditions of each particular transcript or region as a line between the box violins. To ensure accurate comparison, both the control data and the salt stress data were normalized by the same scale (See Methods). For whole transcripts, we observe a small but significant decrease in the average mean transcript reactivity in shoot following salt stress (p=1.80e^−10^; Figure 2A, Tables 1, S.3.a), while in root we observe the opposite trend, with an appreciable increase in the average mean reactivity in whole transcripts following salt stress (p<2.20e^−16^; Fig. 2E, Tables 1, S.3.a).

**Figure 2.**
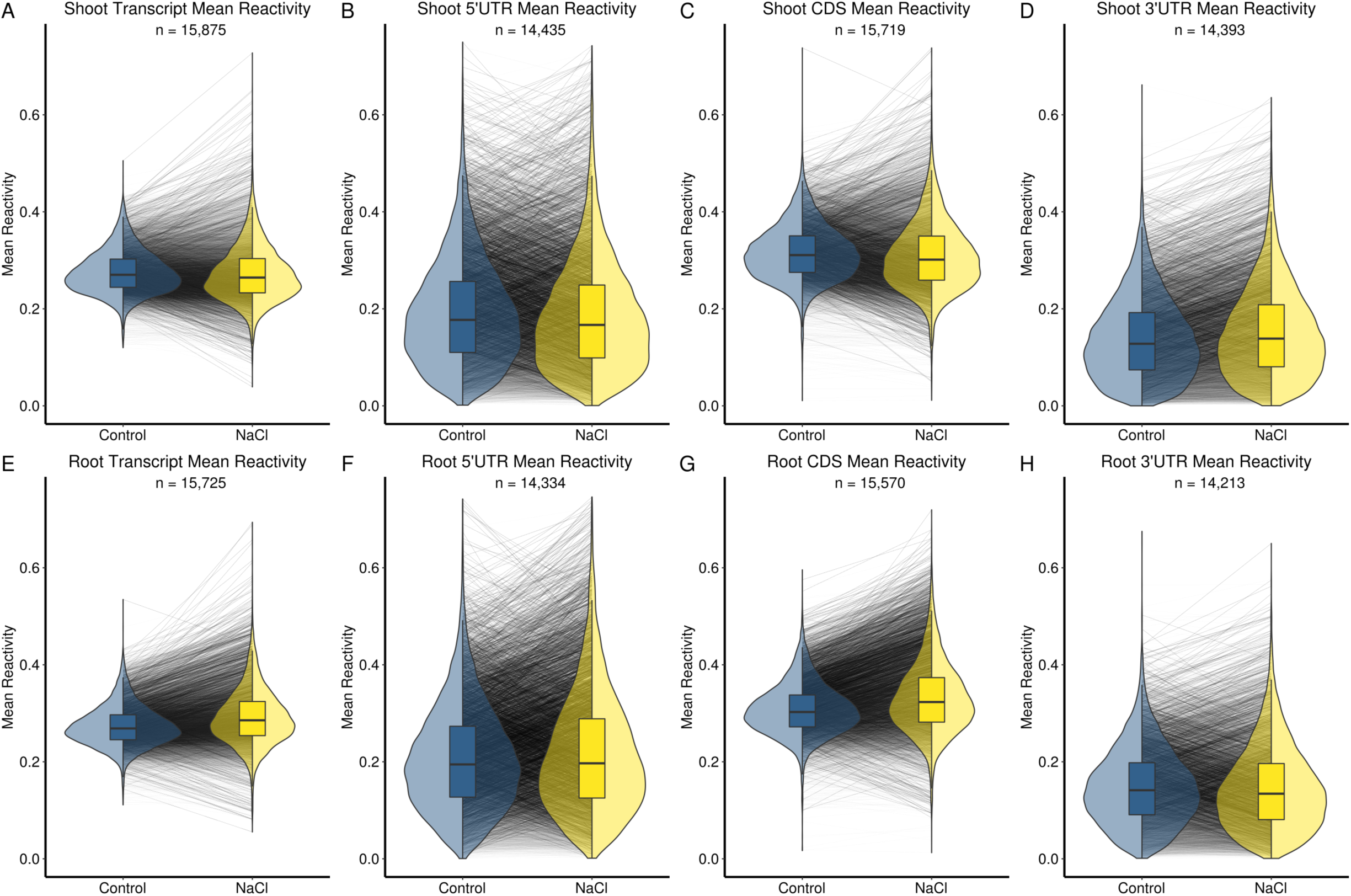
Salt Stress Increases Average Mean Reactivity in Root, but Decreases it in Shoot. distributions of mean reactivity are plotted for both control (blue) and NaCl-treated (yellow) transcripts and transcript regions in both shoot (top row) and root (bottom row). Changes in mean reactivity between conditions of each transcript or region are shown by lines between the distributions. A statistical summary of these results can be found in Tables 1 and S.3.a.

**Table 1.** Summary of Reactivity Changes. Average mean reactivities, standard deviations (SD), and t-test results accompanying Figures 2 and 3. A more complete version of this table can be found in Table S.3.a

| Sample A | Sample B | Category | n | t-test p | Mean A | Mean B | SD A | SD B | r |
| --- | --- | --- | --- | --- | --- | --- | --- | --- | --- |
| Shoot Control | Shoot Salt | transcript | 15,875 | 1.80E-10 | 0.28 | 0.27 | 0.04 | 0.06 | 0.89 |
|  |  | 5'UTR | 14,435 | 8.43E-07 | 0.19 | 0.19 | 0.11 | 0.11 | 0.87 |
|  |  | CDS | 15,719 | 2.20E-16 | 0.32 | 0.31 | 0.06 | 0.07 | 0.88 |
|  |  | 3'UTR | 14,393 | 2.20E-16 | 0.14 | 0.15 | 0.08 | 0.09 | 0.92 |
| Root Control | Root Salt | transcript | 15,725 | 2.20E-16 | 0.27 | 0.29 | 0.04 | 0.06 | 0.88 |
|  |  | 5'UTR | 14,334 | 9.67E-14 | 0.21 | 0.22 | 0.11 | 0.13 | 0.84 |
|  |  | CDS | 15,570 | 2.20E-16 | 0.31 | 0.33 | 0.05 | 0.07 | 0.87 |
|  |  | 3'UTR | 14,213 | 1.43E-03 | 0.15 | 0.15 | 0.08 | 0.09 | 0.88 |
| Shoot Control | Root Control | transcript | 13,974 | 2.20E-16 | 0.27 | 0.21 | 0.04 | 0.07 | 0.68 |
|  |  | 5'UTR | 12,861 | 2.20E-16 | 0.19 | 0.16 | 0.11 | 0.10 | 0.67 |
|  |  | CDS | 13,845 | 2.20E-16 | 0.32 | 0.24 | 0.06 | 0.08 | 0.63 |
|  |  | 3'UTR | 12,803 | 2.20E-16 | 0.14 | 0.11 | 0.08 | 0.07 | 0.80 |
| Shoot Salt | Root Salt | transcript | 13,737 | 2.20E-16 | 0.27 | 0.22 | 0.04 | 0.06 | 0.72 |
|  |  | 5'UTR | 12,681 | 2.20E-16 | 0.18 | 0.16 | 0.11 | 0.10 | 0.72 |
|  |  | CDS | 13,607 | 2.20E-16 | 0.31 | 0.25 | 0.06 | 0.08 | 0.69 |
|  |  | 3'UTR | 12,586 | 2.20E-16 | 0.15 | 0.11 | 0.09 | 0.07 | 0.83 |

Studies on yeast and mammalian cells (Kertesz et. al. 2010, Wan et al., 2012), as well as our recent analyses in rice (Su et al., 2018), have indicated that RNA structure is not uniform throughout the length of mRNAs. We accordingly separated our whole transcript reactivity data into transcript regions (5’UTR, CDS, 3’UTR) for shoot (Figure 2B, 2C, 2D) and root (Figure 2F, 2G, 2H). The average mean reactivity of 5’UTRs marginally decreases in response to NaCl in shoot (p=8.43e^−7^), whereas in root, 5’UTR average mean reactivity increases significantly (p<2.20e^−16^). In the coding region (CDS) under salt stress, the average mean reactivity strongly decreases in shoot and again increases in root (p<2.20e^−16^, Figure 2C, 2G). In the 3’UTRs, a small but significant increase in average mean reactivity in salt conditions occurs in shoot, while the 3’UTRs in root decrease in average mean reactivity (p<2.20e^−16^, Figure 2D, 2H) (Table 1, S.3.a). In both tissues, 5’UTRs and 3’UTRs exhibit greater standard deviation in mean reactivity than the CDS in both control and stress conditions (Tables 1, S.3.a; also apparent in Figure 2 by comparison of the violin plots for the three different transcript regions), suggesting that UTR regions are more structurally malleable and responsive to salt stress than CDS regions (Tables 1, S.3.a). These results highlight the potential of UTRs as structural mediators or regulatory elements.

Different transcripts may have similar reactivity means yet differ greatly in how that reactivity is distributed along the transcript, homogeneously or unevenly. This in turn may have biological consequences. Standard deviation and Gini index have been used to describe the unevenness of this distribution (Rouskin et al. 2014). Standard deviation is affected by the mean value of its metric when normalization is used, thus using standard deviation to measure the differences in how reactivity is distributed along transcripts or transcript regions would bias more reactive transcripts (with higher mean reactivity) to have a higher standard deviation. Thus, we instead use the Gini index to describe the evenness of reactivity within a transcript or region. Gini is a metric that measures evenness of a given set of numbers and is bounded by zero to one, with zero indicating perfect evenness (i.e. all reactivity values are equal) and one indicating perfect unevenness (all reactivity is on a single position). We compared the distributions of Gini reactivity under control and salt stress conditions (Figure S.3, Table S.3.b). With salt treatment, we observe a small but significant increase in the average Gini of reactivity in whole transcript (p<2.20e^−16^, shoot and root), 5’UTR (p<2.20e^−16^, shoot and root), and CDS (p<2.20e^−16^ shoot, p=3.39e^−8^ root) (Table S.3.b), showing that reactivity is more stochastically distributed along mRNAs in both tissues following salt stress. Curiously, the 3’UTR significantly decreases in average Gini reactivity in shoot tissue, yet increases in root tissue after salt treatment (p<2.20e^−16^, both tissues). Taken together with the changes in average mean reactivity, in shoot the 3’UTR gains reactivity and partitions reactivity more evenly in salt stress, while in root the opposite occurs where the 3’UTR loses reactivity and partitions reactivity less evenly in salt stress.

### mRNA Structuromes Show Tissue Specificity

Root tissue contains many cell types known to have specific transcriptomic responses to salt stress in Arabidopsis (Dinneny et al., 2008) and other species (Zhang et al., 2015). Despite the caveat that we could not separate the various cell types’ structuromes within each of our tissues due to the mRNA amount required per each of our 24 libraries, comparing the composite shoot and root structuromes against each other, both in control and salt stress conditions, seemed an interesting avenue of research. For mRNAs present in the structuromes of both tissues, we compared the distributions of mean reactivity (Figure 3, Tables 1, S.3.a) for whole transcript (Figures 3A, 3E) and each transcript region for our control (Figures 3B, 3C, 3D) and salt stress (Figures 3F, 3G, 3H) data in both shoot (left violin, blue) and root (right violin, yellow) tissues.

**Figure 3.**
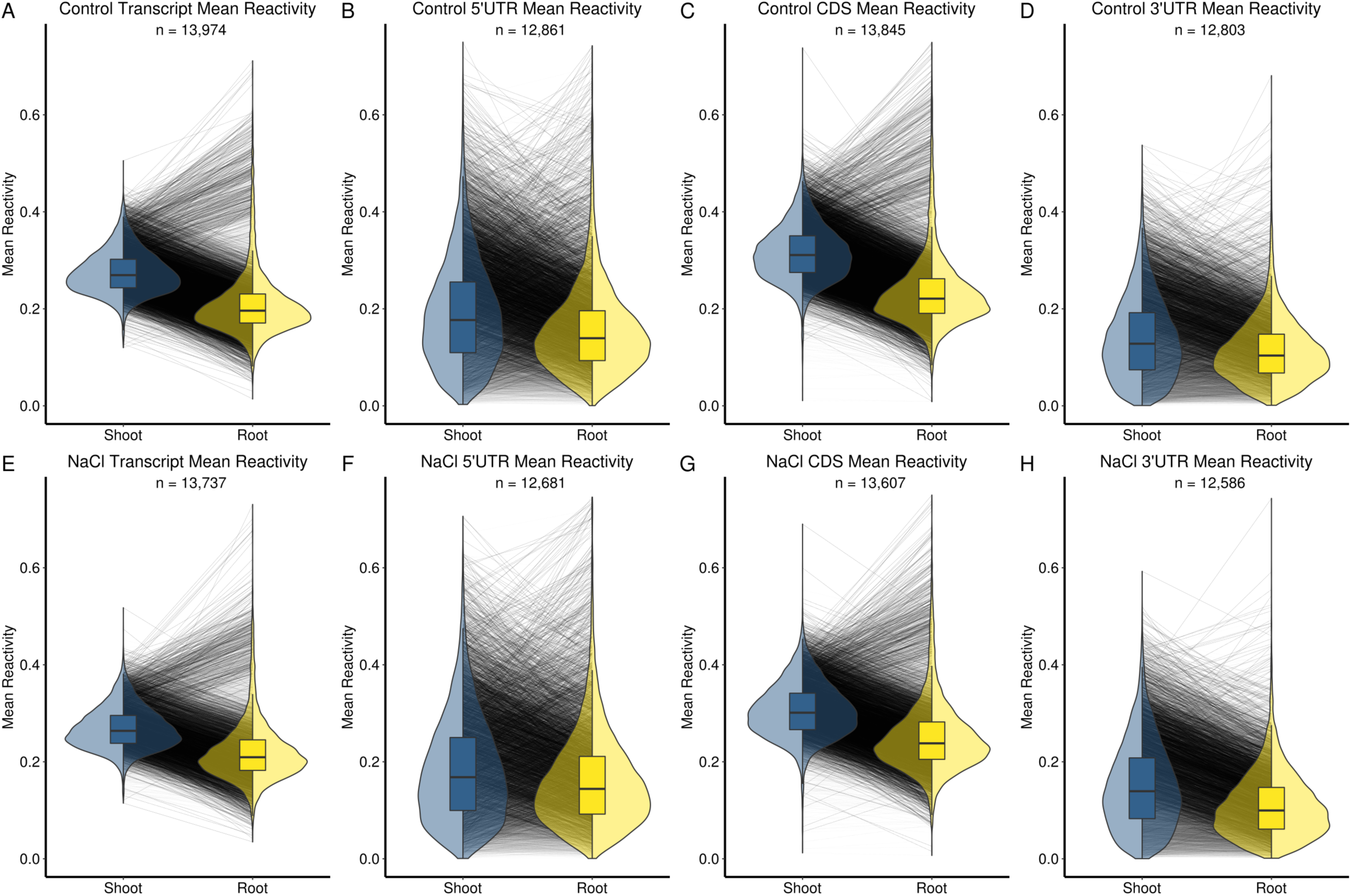
The Effect of Tissue Type and Salt Stress on Arabidopsis Inter-Tissue Transcript Mean Reactivity. Distributions of mean reactivity are plotted for both shoot (blue) and root (yellow) transcripts and transcript regions in both control (top row) and NaCl-treatment (bottom row) conditions. Changes in mean reactivity between conditions of each transcript or region are shown by lines between the distributions. A statistical summary of these results can be found in Tables 1 and S.3.a..

In both control and salt stress conditions, we found a strong trend wherein the global average of the mean reactivity of whole transcripts, and of each transcript region, is distinctly lower in root compared to shoot (p<2.20e^−16^, all regions) (Figure 3, Table 1, S.3.a). The differences between the global averages of shoot and root persist under salt stress for whole transcript, 5’UTR, and CDS regions but become smaller, because the shoot transcriptome generally loses DMS reactivity while the root transcriptome gains reactivity under salt stress, pushing the average means closer. (Figure 3 compare blue to yellow, Tables 1, S.3.a). The 3’UTR behaves differently: in salt, root 3’UTR loses reactivity and shoot 3’UTR gains reactivity, therefore the difference between the average mean reactivities for 3’ UTRs is greater in salt stress. To elaborate on this concept of relatedness, we correlated the mean reactivities of whole transcript and each region between tissues in both conditions. We found that every inter-tissue correlation had a stronger correlation coefficient in salt stress than in control (Table S.3.a); thus, transcript and transcript region mean reactivities are more similar between tissues in salt stress than in control conditions. Extending this line of thinking, we compared all of the intra-tissue (control vs. salt stress) correlations to the inter-tissue correlations (shoot vs. root) (Table S.3.a). These results showed that control and salt-stressed structuromes within a tissue are more related than are the shoot and root structuromes within a condition. In other words, tissue is a stronger determinant of the structurome than salt stress, although both are significant. As an additional measure of difference between the structuromes of shoot and root, we again evaluated changes in the Gini index of reactivity between tissues (Figure S.4, Table S.3.b).

Within each intra-tissue comparison, we divided mRNAs into two categories: (1) those that are found only in shoot or only in root, approximating transcripts that are tissue-specific (unique), and (2) those that are present in both tissues (shared). Then, in each tissue, we compared a variety of metrics between unique and shared transcripts (Table S.4). The absolute value of mean salt-induced change in reactivity was higher in both sets of unique transcripts compared to their respective shared sets, suggesting greater dynamism in the former. Shoot unique transcripts showed 23% more change (p<2.20e^−16^) and root unique transcripts showed 12% more change (p=6.91e^−06^) than shared transcripts.

### ANOVA Confirms the Influence of Tissue-Type on the RNA Structurome

We used four two-way ANOVAs to assess data from both tissues and conditions simultaneously (see Methods, Figure S.5 and Table S.5) rather than comparing just two data sets at a time. The strongest determinants of mean reactivity were tissue and the interaction of treatment and tissue, rather than treatment alone. Thus tissue appears to have an overall stronger influence on RNA structure than salinity stress, as also concluded above. This reinforces the trend wherein the intra-tissue (control vs. salt stress) correlations were stronger than the inter-tissue (root vs. shoot) correlations, and suggests that the structurome is particularly shaped by tissue identity. When ‘region’ (i.e CDS or UTR) was included as a factor in three-way ANOVA, it was also a stronger determinant than treatment.

### Salinity-Induced DMS Reactivity Changes Inversely Correlate with Salinity-Induced mRNA Abundance Changes

We recently reported a significant inverse relationship between change in transcript DMS reactivity and change in transcript abundance in rice seedlings after 10 min. of heat shock (Su et al., 2018). We hypothesized that this might be a general phenomenon underlying abiotic stress regulation of transcriptomes in multiple species and under multiple stresses. Accordingly, we compared the changes of mean reactivity (Δ reactivity) to the changes of abundance (Δ abundance) calculated from respective pools of -DMS libraries and expressed as log_2_ (TPM) (Transcripts per kilobase-Million) between control and salt conditions for whole transcript and each transcript region in both shoot and root (Figure 4, Tables 3A, S.6.a). We found a striking negative correlation between Δ reactivity and Δ abundance (p<2.20e^−16^, both tissues, all regions). In each tissue, whole transcript and CDS showed the strongest negative correlation, followed by 5’UTR and lastly 3’UTR. The relationship was stronger in each shoot comparison than in each root comparison (e.g. whole transcript shoot r= −0.73, root r= −0.58). Correlations were also calculated for subsets consisting of the shared and unique (tissue-specific) transcripts (Tables 2B, S.6.b). In each case tissue-specific transcripts exhibited stronger correlations than tissue-shared transcripts (see Discussion).

**Figure 4.**
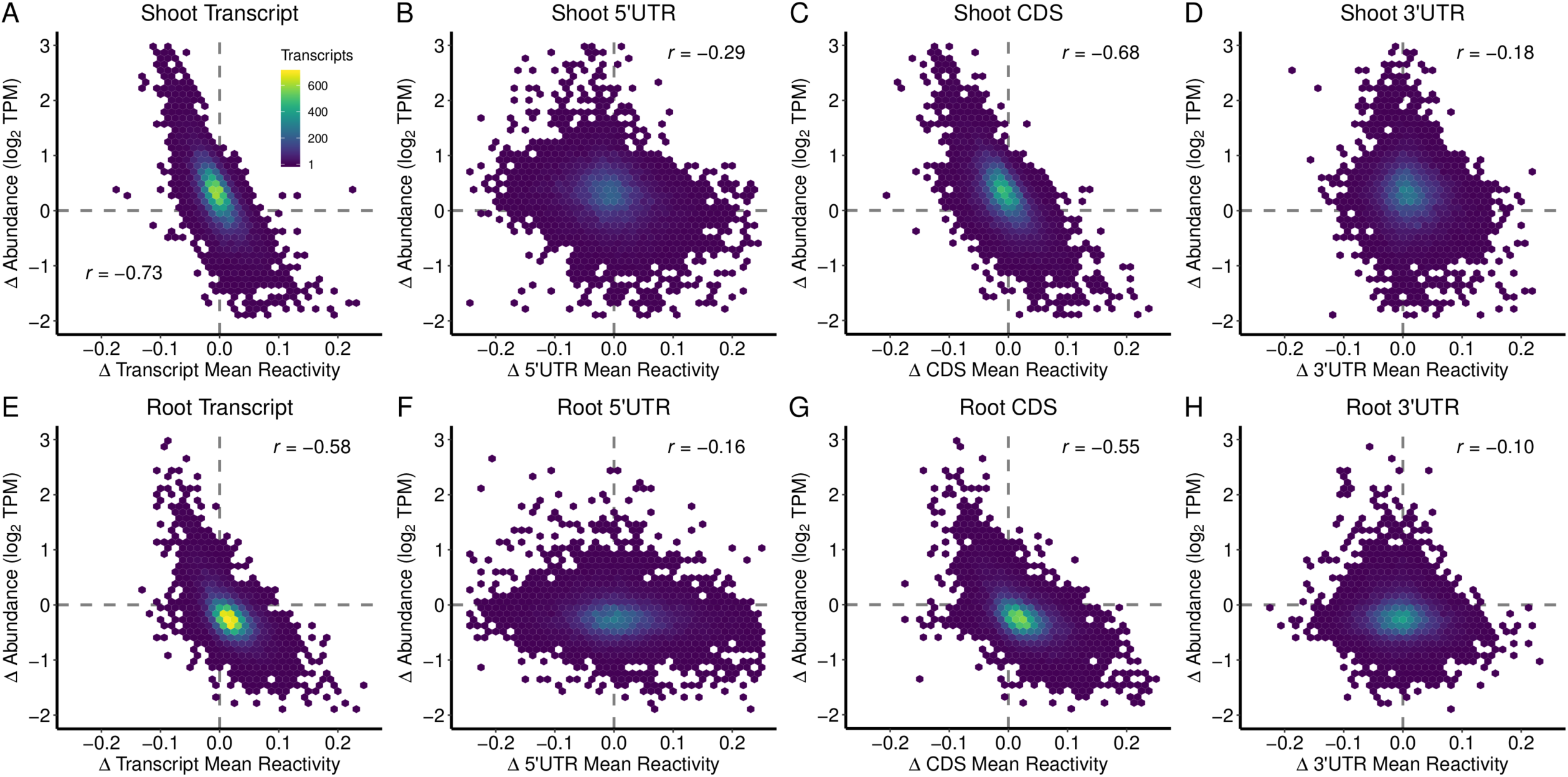
Change in Reactivity is Inversely Correlated with Change in Abundance. For each transcript and transcript region, we assessed the relationship between change in mean reactivity (Δ mean reactivity, NaCl-treatment mean reactivity - control mean reactivity, x-axis) to the change in relative abundance (Δ abundance, log2(TPM NaCl) - log2(TPM control), y-axis) between control and NaCl treatment in shoot and root tissues. This revealed a strong inverse correlation (Tables 2A, S.6.a).

Based on *in vitro* analyses of the Arabidopsis RNA structurome based on digestion with single-stranded and double-stranded nucleases, Gregory and co-workers (Li et al. 2012) reported a direct (positive) relationship between abundance and RNA single-strandedness, seemingly in contradiction to our observations of an inverse (negative) relationship governing *changes* in these two parameters. We resolved this apparent paradox as follows. To comprehensively assess this trend in a manner as close as technically possible to Li et al., we re-calculated all reactivities for every transcript and transcript region in each condition and treatment individually (Tables R.1-R.16), i.e. one structurome at a time. This allowed a greater number of transcripts per individual structurome to investigate this relationship because mutual coverage between conditions did not have to be considered. Additionally, using each structurome’s internal normalization scale mitigates some dispersion of reactivity values. When we assess these datasets for the relationship between whole transcript mean reactivity and transcript abundance, we find a strong positive correlation (r ≥ 0.75, p<2.20e^−16^, Figures S.6-S.9, Table S.6.c) between mean DMS reactivity and transcript abundance as anticipated from the studies of Li et al. (2012). This correlation appears driven primarily by the reactivity of the CDS, as it was only weakly observed for either 5’ or 3’ UTRs (Figure S.6, Table S.6.c). Having confirmed the trend in individual structuromes, we also confirmed it in our contrasted structuromes (Figures 5A, 5D, Table S.6.d), which were used to investigate how this relationship is affected by changes between conditions. In response to salt stress, highly abundant (i.e. high reactivity) transcripts preferentially further increase in reactivity and decrease in abundance. Low abundance (i.e. low reactivity) transcripts preferentially further decrease in reactivity and increase in abundance following salt stress (Figures 5B, 5C, 5E, 5F, S.7, Table S.6.e) (see Discussion).

**Figure 5.**
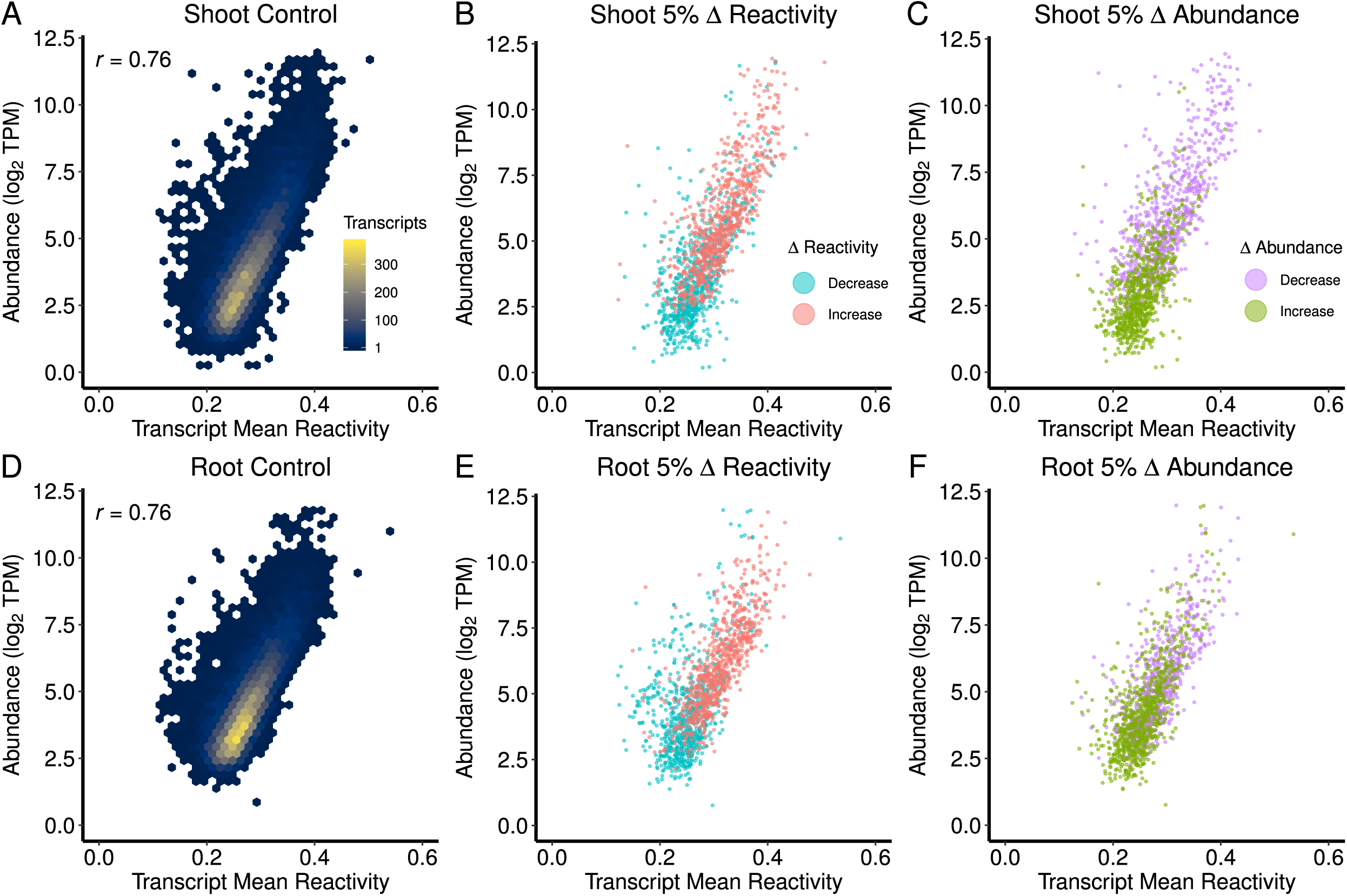
Relationship between Reactivity and Abundance and its Modulation by Salt Stress. A,D) Transcript control mean reactivity (x-axis) is positively correlated with transcript control abundance (y-axis). Data are for transcripts resolved in both control and salt-stress conditions within both Shoot (A) and Root (D) (Table S.6.d). B,E) Transcripts within the top and bottom 5% of salt stress-induced reactivity changes are subset, showing that high abundance transcripts in control conditions tend to increase in mean reactivity under salt stress, while low abundance transcripts tend to decrease in mean reactivity. C,F) Transcripts within the top and bottom 5% of salt stress-induced abundance changes are subset, showing that high abundance transcripts in control conditions tend to lose abundance under salt stress, while low abundance transcripts tend to increase in abundance (Table S.6.e).

### Concordancy Strengthens the Inverse Relationship between Reactivity Changes and Abundance Changes

The observation that the strongest inverse correlation of Δ reactivity to Δ abundance occurred in the CDS (Figure 4, Tables 2A, S.6.a) was unexpected given that previous analyses appear to implicate UTRs as more structurally dynamic and directly or indirectly tied to mRNA decay mechanisms (Su et al., 2018; Wan et al., 2012). We accordingly classified our UTRs as concordant (share the same sign of Δ reactivity in response to salt stress) or discordant (do not share the same sign) with respect to the other transcript regions. In both shoot and root, we compared the Δ reactivity of each transcript region against the Δ reactivity of each other region in the transcript (5’UTR:CDS, 3’UTR:CDS, 5’UTR:3’UTR) (Figures S.8 and S.9 (left columns), Table S.7.a), finding no overall relationship between the reactivity change between different transcript regions. Thus, concordancy between regions of a given transcript is uncommon.

We parsed out transcripts where all three transcript regions were concordant in their salt-induced reactivity changes (Figure 6, Tables 2C, S.7.c). Out of the 15,875 shoot and 15,725 root transcripts resolved, 3,155 and 3,175 respectively were fully concordant (~20%), leaving 12,720 and 12,550 (~80%) discordant transcripts (meaning two of the three regions have different changes in reactivity). Strikingly, whole transcripts and each transcript region show their most significant respective Δ reactivity to Δ abundance correlation value when all regions are concordant in their direction of salt-induced change in reactivity, and this relationship is observed in both shoot and root tissues (p<2.20e^−16^, both tissues, all regions): *concordant* whole transcripts shoot r = –0.82, root r = –0.65 (Figure 6), contrasted to *all* transcripts whole transcript r = –0.73 shoot, r = –0.58 root (Figure 4). Correlation values are generally lower for root than for shoot. Of the 3,155 and 3,175 fully concordant transcripts in shoot and root respectively, 2,596 (~82%) and 2,688 (~84%) were present in both tissues, but only 628 (~23%) of these shared transcripts were fully concordant in both tissues, suggesting that tissue-type is a strong determinant of a transcript’s concordancy. Within this set of 628 concordant shared transcripts, only 321 (~51%) also showed the same direction of concordancy change in both tissues (transcript lists S.8), further pointing toward tissue-specificity of the concordancy relationship.

**Figure 6.**
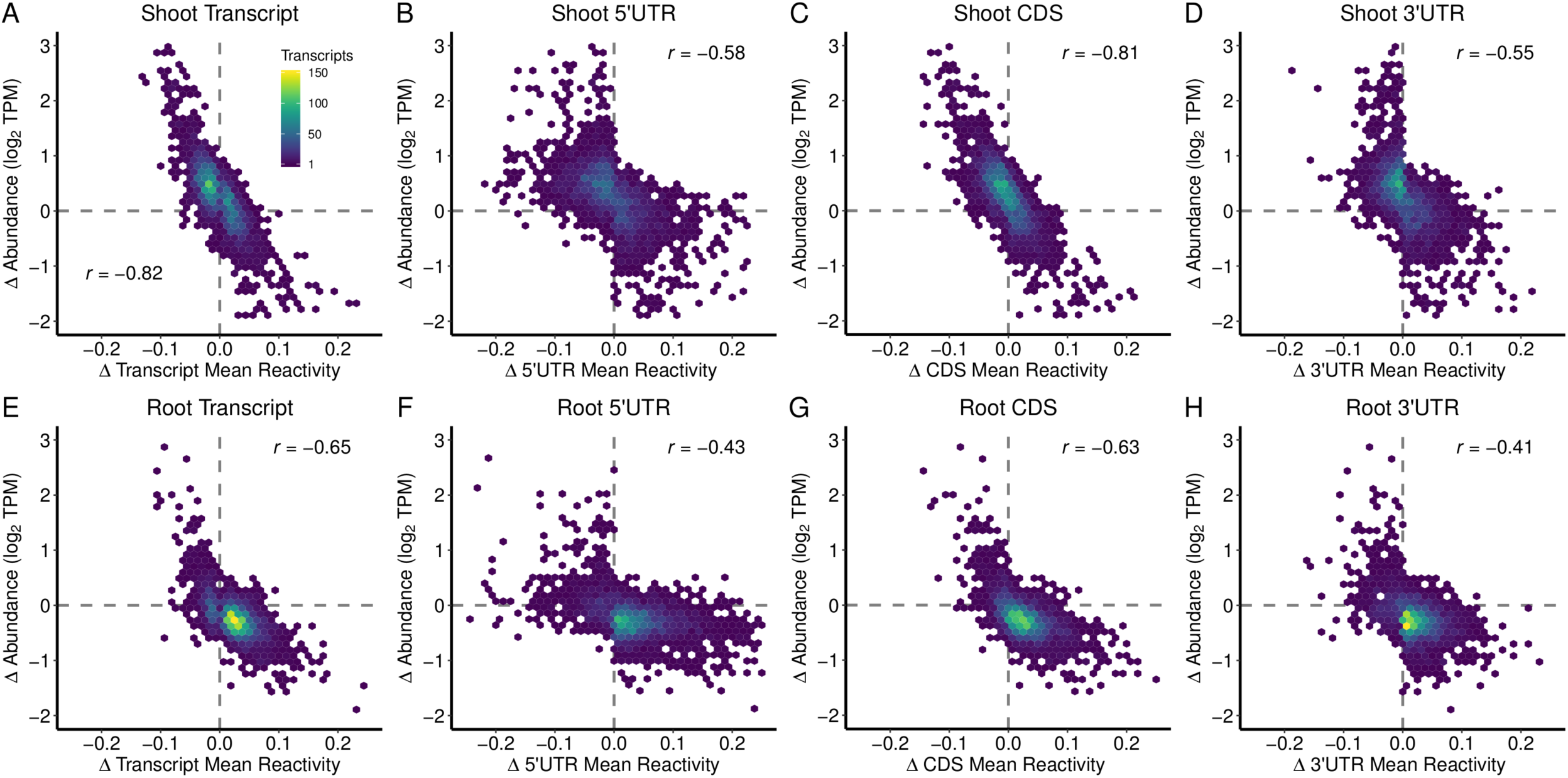
Concordant Reactivity Change has the Strongest Influence on Abundance Change. Transcripts that share the same sign of Δ mean reactivity between 5’UTR, CDS, and 3’UTR regions were subset as concordant. For each transcript and transcript region of these fully concordant transcripts, we assessed the relationship between the change in mean reactivity (Δ mean reactivity, NaCl mean reactivity - control mean reactivity, x-axis) and the change in relative abundance (Δ abundance, log2(TPM NaCl) - log2(TPM control), y-axis) between control and NaCl-treatment in shoot and root tissues, revealing a strong enhancement of the inverse correlation found among all transcripts and regions (Figure 4, Tables 2A, S.6.a) (Tables 2C, S.7.c for all concordant tests and numbers).

Next, we assessed the strength of the Δ reactivity versus Δ abundance relationship between every two pairs of transcript regions (5’UTR:CDS, 3’UTR:CDS, 5’UTR:3’UTR), sub-grouping them into concordant and discordant categories. Concordancy was defined as transcript regions sharing the direction of reactivity change between conditions and discordancy as not sharing the direction of reactivity change between conditions. This leads to 24 different combinations, which are presented as Δ abundance versus Δ reactivity plots in Figures S.8 (shoot), S.9 (root), with all tests and statistics available in Table S.7.b. Regions concordant in Δ reactivity with at least one other region have a stronger inverse Δ reactivity-per-region to Δ abundance-per-transcript relationship than discordant regions. For instance, transcripts with 5’UTRs that are concordant in the direction of reactivity change with their respective CDS have a stronger relationship to abundance change (–0.51 shoot, –0.39 root, p<2.20e^−16^), than transcripts with 5’UTRs that are discordant in the direction of reactivity change with their respective CDS and even have a slight direct correlation (+0.15 shoot, +0.21 root, p<2.20e^−16^). These results suggest that interplay between the structuredness of two different transcript regions (5’UTR, CDS, 3’UTR) can dynamically contribute to the Δ reactivity versus Δ abundance relationship.

### Gene Ontology Analyses

We performed 36 individual GO analyses via agriGOv2 (Tian et al. 2017), testing for the enrichment of categories that change in each contrast (full index of analyses in Table G.0). For our intra-tissue contrasts, we examined the top and bottom 5% of increased and decreased mean reactivity between control and salt stress conditions for whole transcript and each transcript region in both tissues (2×2×4 = 16 intra-tissue analyses, Tables G.1 to G.16). For our inter-tissue contrasts, we examined the top and bottom 5% of increased and decreased mean reactivity between tissues for whole transcript and each transcript region in both conditions (2×2×4 = 16 inter-tissue analyses, Tables G.17 to G.32). For GO analyses on concordant transcripts, we subset our transcripts into those that increased in abundance and lost reactivity (concordant protection) and those that decreased in abundance and increased in reactivity (concordant exposure) between conditions in both tissues (2×2 = 4 concordant analyses, Tables G.33 to G.36), using agriGOv2 (Tian et al. 2017). When submitting queries to agriGOv2, we used genome locus rather than transcript such that genes with multiple transcripts present in our data did not gain over-representation.

We present some of these findings in Figures 7 and 8. We find that increases in reactivity during salt stress in whole transcript and each transcript region are categorically enriched in genes related to photosynthesis in shoot, and that such increases in both tissues are enriched in response to oxidative stress (Figure 7A). For transcripts and transcript regions that decrease in reactivity during salt stress, both tissues show enrichment in many stress response categories (Figure 7C). Notably, in both tissues, we find enrichment in response to salt stress and response to osmotic stress, as well as response to water deprivation in concordant loss of reactivity/gain in abundance transcripts (Figure 8). In shoot, genes related to photosynthesis lose abundance and are concordant in gain of reactivity across all transcript regions (Figure 8), consistent with only aerial tissues performing photosynthesis and exhibiting reduced photosynthesis under stress.

**Figure 7.**
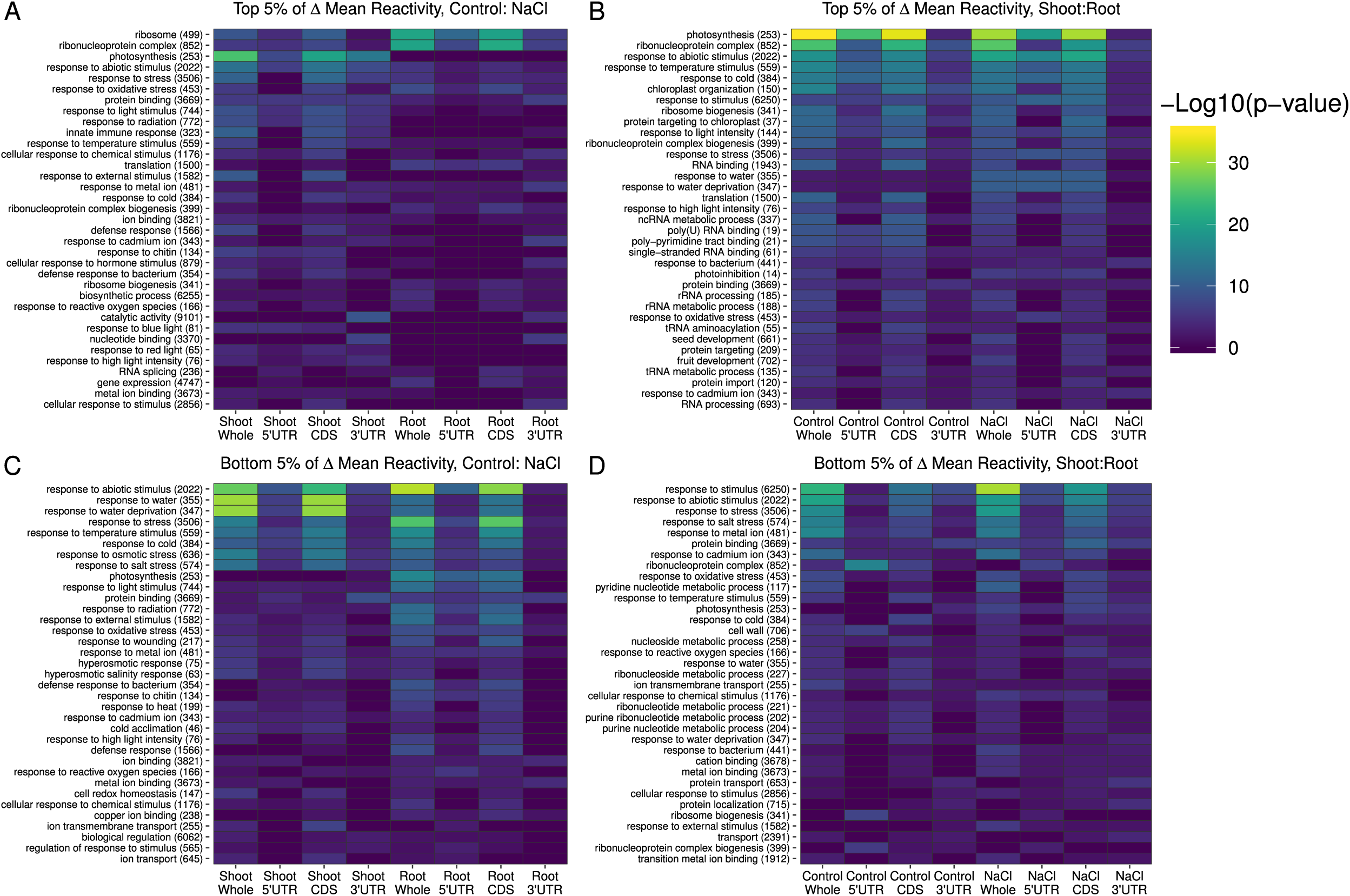
Gene Ontology Analyses Reveal Enrichment of Stress-related Transcripts. Selected results from multiple GO analyses (via agriGOv2, Tian et al. 2017) are combined and displayed as a heat map, where color indicates the significance of enrichment within an analysis. A,C) Transcripts and transcript regions within the top 5% of reactivity increases or decreases after salt stress. B,D) Transcripts and transcript regions within the top 5% of reactivity differences between tissues. The top 5% of Δ shoot:root indicates transcripts that are more reactive in root, and the bottom 5% Δ shoot:root indicates transcripts that are less reactive in root. The full results of these GO analyses are found in Tables G.1-G.32.

**Figure 8.**
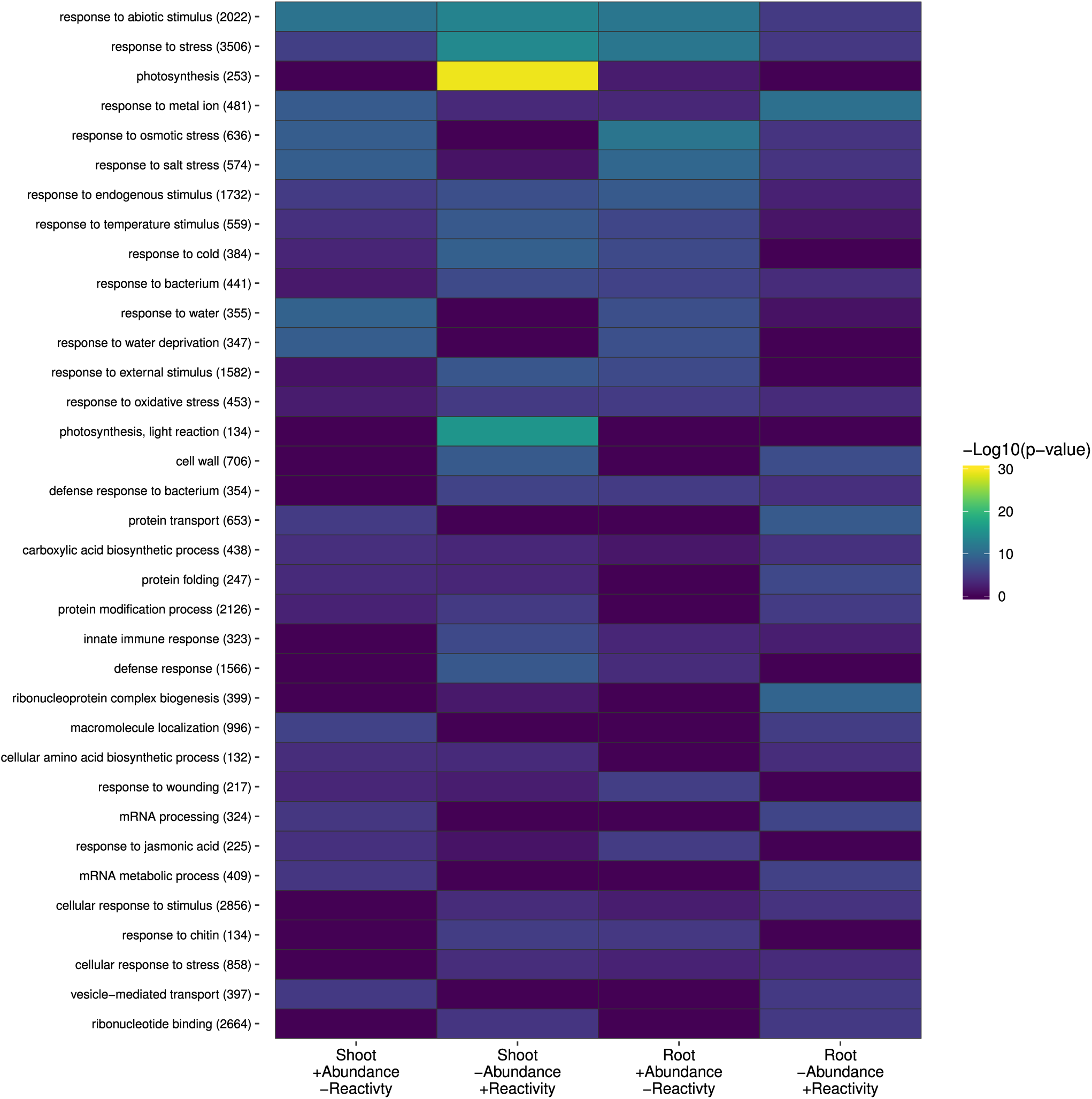
Concordant Transcripts Are Enriched in Stress-Related Gene Functions. Concordant transcripts from both shoot and root were separated into those that increased in abundance and decreased in mean reactivity (+abundance/-reactivity) and those that decreased in abundance and gained in mean reactivity (-abundance/+reactivity) after salt stress. These subsets were used in a series of GO analyses using agriGOv2 (Tian et al. 2017) and summarized as a heatmap; the full results can be found in tables G.33-G.36. The set of (+abundance/-reactivity) transcripts shows enrichment in categories pertinent to salt stress réponse. Conversely, the (-abundance/+reactivity) set of transcripts show categorical of functions not specifically associated with salt stress, suggesting concerted down regulatory action attenuates the abundance of these transcripts. Concordant change of all three transcript regions therefore may particularly modulate stress-induced abundance changes.

### Changes in Predicted RNA Structure

To approximate actual structural effects induced by salt, we used the DMS reactivity profiles as restraints to guide folding (Tack et al. 2018; Ding et al., 2014; Deigan et al. 2009) of transcripts (see Methods). There were 15,708 mRNAs in shoot and 15,557 mRNAs in root with sufficient data to generate structures in both control and salt conditions. For each pair of structures (control and salt) in a tissue, we calculated PPV (Positive Predictive Value) (Tables F.1-F.8): the fraction of base pairs shared between the same transcript in the same tissue under control versus salt stress conditions. The distributions of PPV are summarized in Figure 9A, showing that transcripts in root tissue have a lower mean PPV, indicative of fewer shared base pairs between control and salt-stressed conditions as compared to shoot. This result is consistent with the bigger gain in DMS reactivity in root than loss in reactivity in shoot (see Figure 3 and Table 1).

**Figure 9.**
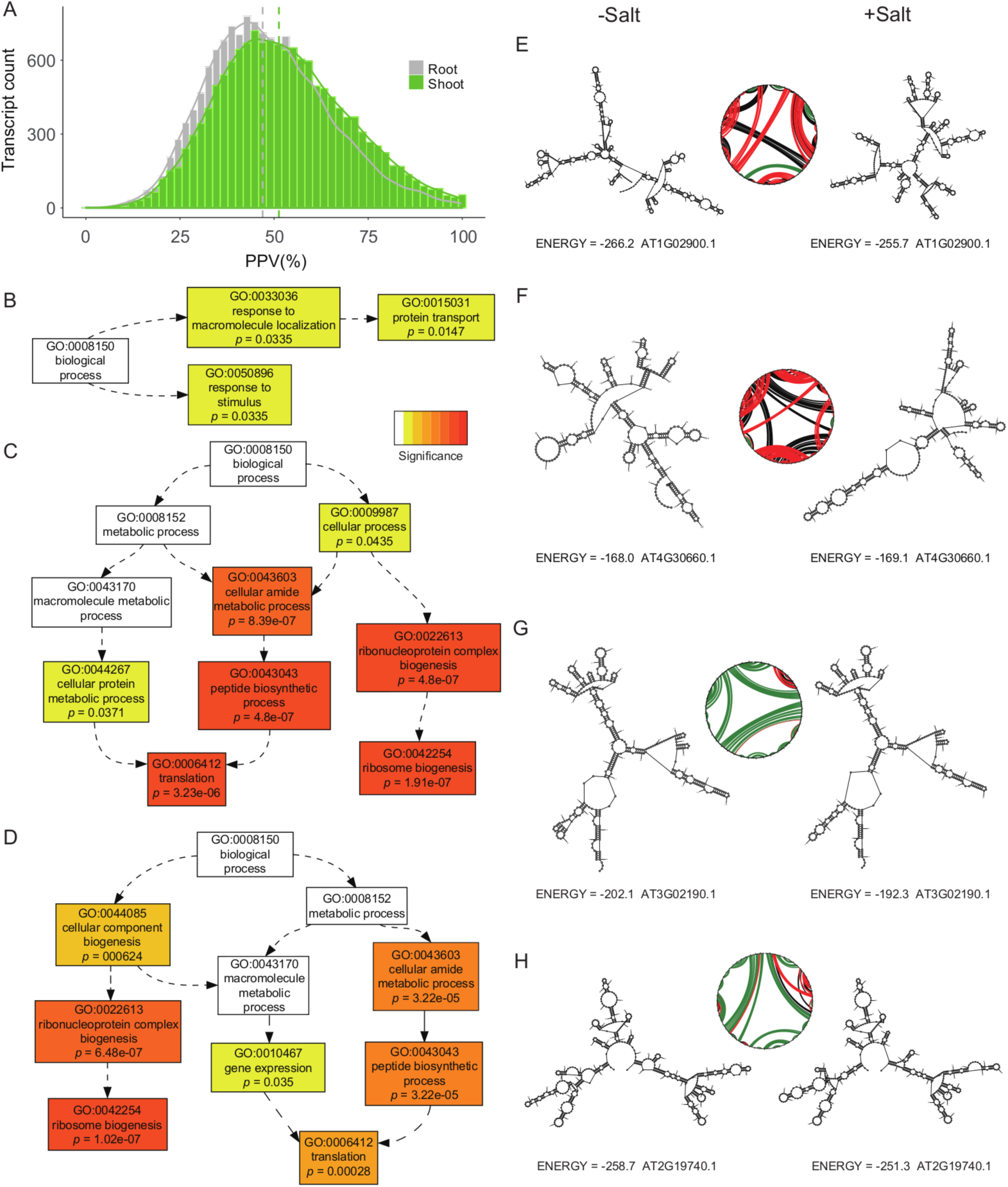
Folding Analyses Suggest Core Function of RNA Structure in Salt Stress Response. A) PPV distribution of 15,708 mRNAs in shoot (green) and 15,557 mRNAs in root (grey). PPV values quantify the structural changes between salt stress and control condition. Dashed lines indicate the mean PPV in root (grey; mean: 46.96) and shoot (green; mean: 51.30). PPV is the fraction of base-pairs that appear in two structures of the same RNA so is bounded by 0 and 1, with 1 indicating identical structures. B) GO categories “response to stimulus” and “protein transport” are overrepresented in the 5% of the transcripts with lowest PPV in root (but not in shoot). C) Transcripts involved in translation and ribosome biogenesis were highly enriched in the 5% of transcripts with highest PPV in shoot. D) Transcripts involved in translation and ribosome biogenesis were also highly overrepresented in the 5% of transcripts with highest PPV in root. E) An example of a low PPV transcript (AT1G02900.1, rapid alkalinization factor 1 (RALF1), PPV = 22.7) showing obvious structural changes between control condition (left) and salt stress (right). F) Another example of a low PPV transcript (AT4G30660.1, Low temperature and salt responsive protein family gene, PPV = 0) with dramatic structural change between control condition (left) and salt stress (right). G) An example of a high PPV transcript (AT3G02190.1, 60S ribosomal protein L39 (RPL39B), PPV = 87.79) with similar structure between control and salt stress. H) Another example of a high PPV transcript (AT2G19740.1, 60S ribosomal protein L31 (RPL31A), PPV = 97.38) with similar structure between conditions. For E, F, G, H, all structures were generated from root data. Circle plots provide an abstraction of the base pairing (Reuter and Matthews, 2010). Nucleotides are arranged in sequence order around the circle. Black lines link bases that are paired only in control; red lines link bases that are paired only in salt stress, green lines link bases that are paired in both conditions. All PPV statistics logged in supplemental tables F.1-F.8.

We took mRNAs in the 5% bottom and top tails of the PPV distributions and submitted them to AgriGO v2.0 (Tian et al. 2017). These GO analyses revealed that transcripts with low PPV (i.e. that show large salt stress-induced changes in predicted structure) were enriched in genes associated with response to stimulus (GO 0050896; Figure 9B) and protein transport (GO 0015031) in the root. Transcripts with high PPV (i.e. exhibiting similar predicted structure between control and salt stress conditions) were enriched in gene functions associated with translation (GO 0006412) and ribosome biogenesis (GO 0042254). Enrichment of these GO categories was observed in both root and shoot samples (Figure 9C,D). Figure 9E, 9F provides examples of specific transcripts from the low PPV set and Figure 9G, 9H provides examples of specific transcripts from the high PPV set, presenting both predicted secondary structures and an abstraction of their base-pairing in circle plots (Reuter and Matthews, 2010). Salt-induced structural changes are clearly evident in the structures in Figure 9E,9F, while structures in Figure 9G, 9H are highly similar in control and salt conditions.

## Discussion

RNA performs many distinct roles in the cell and has the ability to switch conformations in differing chemical environments. Examples of changes in RNA structure that confer molecular function include RNAs functioning as small molecule (Mandal and Breaker 2004; Wachter et al. 2007) or thermal sensors (Narberhaus et al., 2006) and assisting in translational control (Parker and Sheth, 2007). We and others have recently developed ways to probe RNA structure *in vivo* and genome-wide, implicating functional roles of RNA structure in splicing, polyadenylation, and stress response (Ding et al. 2014; Rouskin et al. 2014; Wan et al. 2014), and suggesting the conservation of structure between phylogenetically distant plant species (Deng et al., 2018). Analyzing the rice *in vivo* mRNA structurome post heat-shock, we recently revealed a dynamic interplay between mRNA structure and mRNA abundance on a genome-wide scale (Su et al., 2018). Here we build on those findings with the initial investigation of tissue-specific structural differences, as well as RNA structural changes invoked by a salinity stress treatment.

### Structuromes are Tissue-Specific

While RNA-seq methods have revolutionized the study of tissue and cell-specific transcriptomes (Pennisi, 2018), to our knowledge the possibility of tissue specificity in mRNA structure has never been assessed. We directly compared the shoot and root structuromes in stressed and unstressed conditions. We discovered that tissue has a stronger influence on the structurome than salt stress; that is, there is a greater difference between the shoot and root structuromes (Figure 4) than between salt stress and control condition structuromes within either one of the tissues (Figure 3). Our ANOVA analyses (Figure S.5, Table S.5) parse out the relative contributions of tissue, stress, and mRNA region to explain DMS reactivity in a broad sense, providing rigorous statistical significance of tissue as the strongest determinant of reactivity. If future technique development can reduce the quantity of RNA required per each custom Structure-seq library, it would be of great interest to dissect these tissue-specific signatures by creating and analyzing Structure-seq libraries from the component cell types of each tissue (Dinney et al. 2008). In particular, we found that transcripts that were not detected in both tissues, i.e. that were tissue-specific in our datasets, had greater mean reactivity change between control and salt conditions (Table S.7.a) and stronger Δ reactivty to Δ abundance correlations (Table S.7.b) than transcripts that were shared between tissues, hinting that transcripts specialized to a tissue have less adaptive constraints on the malleability of their structures.

### Shoot and Root Structuromes Converge in Salt Stress

The shoot and root structuromes become more similar to each other in salt stress relative to control conditions, with the shoot transcriptome generally losing reactivity and the root transcriptome gaining reactivity (Figure 3, Tables 1, S.3.a). The observed trends suggest that both tissues may be changing toward a common stress response optimum. We surmise that some of this trend may be explained by NaCl-induced loss of the high K^+^ levels present in the root tissue under control conditions (Figure 1, mirrored in Yu and Assmann 2015) leading to nearly the same total K^+^ levels in both tissues.

Generally speaking, monovalent cations promote RNA folding and disfavor RNA-protein interaction (Bloomfield et al., 2000). In shoot measured total monovalent ion concentration increases from about 35 to 60 mg/dry weight (g), while in root it decreases from about 80 to a similar final value of 65 (Figure 1, Table S.1). Although available techniques do not query where these ions reside within the cell, this trend is nonetheless consistent with shoot becoming less DMS reactive and root becoming more DMS reactive upon salt stress, assuming effects are dominated by RNA folding rather than protein unbinding.

It should be noted, however, that increasing levels of monovalent cations can *disfavor* RNA folding when adequate divalent ions are present (Manning, 1972; Heilman-Miller et al., 2001); what scenario is operative *in vivo* is not clear, given the complexity of Mg^2+^ and other ion concentrations *in vivo* (see for example Figure S.1, panel F, showing decreased Mg^2+^ content following salt stress), and the presence of a changing complex mixture of osmolytes (see for example Figure 1C). Nonetheless, it is clear that ionic conditions are converging between shoot and root and thus the biophysics, no matter how complex, should also be converging. In other words, the convergence of total ion content per dry weight in the two tissues during stress is consistent with the convergence of DMS reactivity in the two tissues during stress. Again, these ionic changes do not support the hypothesis that protein protection is highly explanatory of our observations.

### Reactivity Change Influences Abundance Change

We recently reported the first observation of a negative correlation between DMS reactivity change and mRNA abundance change, occurring in rice seedlings in response to just 10 min of heat shock (Su et al., 2018). Further analyses suggested that degradation by the exosome and the 5’ endonuclease XRN4 contribute significantly to rapid loss in transcript abundance following heat shock (Su et al., 2018). Here, we also observe a striking inverse relationship between DMS reactivity changes and transcript abundance changes in Arabidopsis shoot and root tissue following 2 days of salinity stress (Figure 4, Tables 2A, S.6.a). These observations suggest a general principle whereby mRNA unfolding following a stress is causative of decreases in mRNA abundance. While longer term stresses such as that imposed here allows ample time for transcriptional regulation, the results reported here and in Su et al. (2018) highlight the importance of looking beyond transcriptional control alone, and reinforce the importance of RNA structure as an environmental sensor and principal driver that reshapes the transcriptome in response to the environment.

We compared (Figure 10) our salt stress-induced abundance changes in both shoot and root (y-axis) to those reported in a recent RNA-seq analysis from Gregory and co-workers (Anderson et al., 2018) (point color) of Arabidopsis rosettes following a two-week salt stress. Our shoot NaCl-induced abundance changes match the abundance changes that Anderson et al. reported (Figure 10). Moreover, in general, transcripts that lost abundance in the Anderson study gained DMS reactivity in our study, while transcripts that gained abundance in the Anderson study lost DMS reactivity in our study (Figure 10). First, these comparisons support the conclusion that the transcriptome changes we observe typify salinity impacts on the Arabidopsis transcriptome. Second, beyond validating our expression results and the effectiveness of our salt stress treatments, the congruency between these datasets further implicates RNA structure as an essential mechanism that post-transcriptionally regulates mRNA abundance. The Anderson et al. study discovered a role of m6A modification in protecting mRNAs from cleavage under salinity stress, providing one example of post-transcriptional regulation. Nonetheless, upon excluding from our datasets all transcripts with an m6A methylation site detected in the Anderson et al. report, our inverse relationship between change in transcript reactivity and change in transcript abundance is strongly preserved (Figure S.10). This result indicates that structure-related mechanisms, independent of m6A modification, dynamically impact transcript abundance. Future studies will help identify underlying mechanisms.

**Figure 10.**
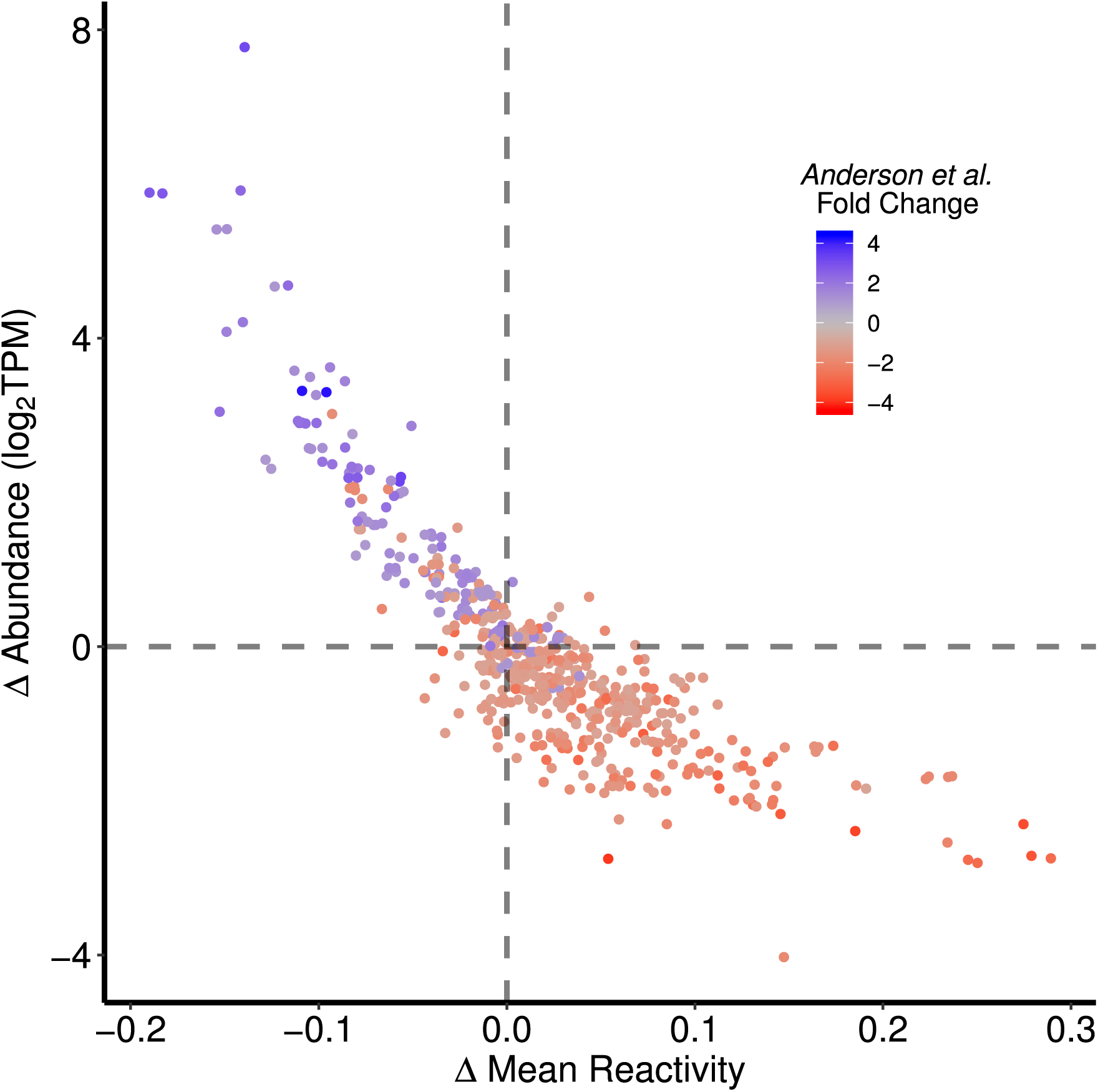
Extended Confirmation of Delta Reactivity Versus Delta Abundance Relationships. Transcripts from our shoot data originating from a gene that had differential expression under salt treatment in Anderson et al. (2018) are plotted. These transcripts are a subset of our whole transcript delta Reactivity versus delta Abundance data in Figure 5. Each dol represents a single transcript colored by its respective gene’s fold change during NaCl treatment extracted from Anderson et al. (2018). An overwhelming majority of NaCl-induced fold changes reported in Anderson et al. (2018) are paralleled in our study, showing parity between the effect of salt treatment on transcript abundance between studies while also highlighting these transcripts’ corresponding changes in reactivity in our study.

We further studied the distribution across the transcriptome of these structurally responsive transcripts (Figure 5, Tables S.6.d, S.6.e). We found a bias wherein abundant transcripts preferentially show concomitant abundance decreases with reactivity increases following salt stress, while rare transcripts preferentially show concomitant abundance increases with reactivity decreases following salt stress (Figure 11). We speculate that the latter constitute specialized stress-related mRNAs that are upregulated in response to salinity, while the former encode everyday metabolic functions. Our GO analyses (see next section) support that the latter are specialized stress-related mRNAs that are upregulated in response to salinity, while the former include transcripts encoding everyday metabolic functions. Thus, this pattern may reflect a repartitioning of resources away from basic metabolism and toward stress amelioration perhaps analogous to growth-defense tradeoffs (Huot et al. 2014).

**Figure 11.**
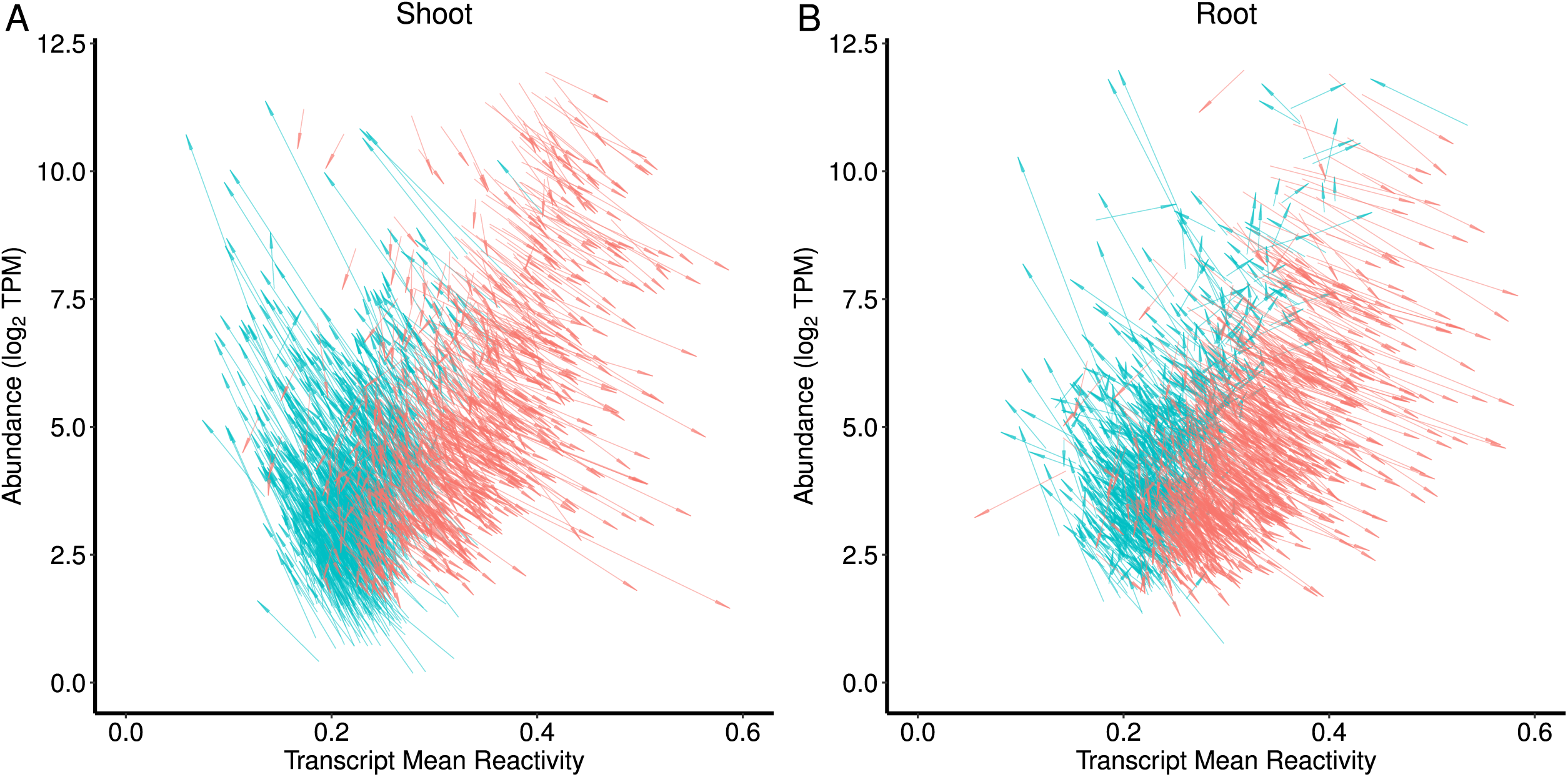
Salt Stress-Induced Reactivity and Abundance Changes as Vectors. For both shoot (A) and root (B), the top 5% most extreme increases (blue) and decreases (red) of transcript abundance from control to salt stressed conditions are plotted as vectors of change. Each vector originates at the (x,y) coordinates of the transcript control values (mean reactivity, log2(TPM)) and ends at the mean reactivity and abundance values the transcript takes on in salt stressed conditions (arrowhead). This further expands on Figures 5 and S.7, distinctly showing the actual vectors of change wherein low abundance transcripts lose reactivity but gain abundance, and high abundance transcripts gain reactivity and lose abundance. In this manner, the transcriptome (abundance) appears reshaped in stress in part by changes in the structurome (reactivity).

### Novel Concordancy Analysis Reveals Synergistic Roles of Transcript Regions

Our previous studies on rice revealed that RNA structure is not uniform between transcript regions, nor are these regions uniformly plastic in response to heat shock (Su et al., 2018). Our present study also reveals particular diversity in the structural responses of the UTR regions compared to the CDS regions following stress, here salinity stress (Figure 2, Tables 1A, S.3.a). Although the CDS shows a particularly strong (inverse) correlation, the structures of the 5’UTR, CDS, and 3’UTR each appear to contribute to regulation of transcript abundance (Figure 4, Tables 3A, S.6.a). Sequence motifs in the 3’UTR that control transcript degradation are known (Rabani et al., 2017), yet little information exists on how structure plays into this. A recent study on mRNA decay in Arabidopsis showed that transcripts with enriched A content and depleted G content in 5’ UTRs exhibited faster decay rates (Sorenson et al. 2018). Given that AU base-pairing is weaker than GC base-pairing, these results may implicate salt-induced 5’ UTR unfolding (increased reactivity) as a mechanism that exposes transcripts to nucleolytic attack. While the CDS region is under selection for peptide sequence and codon efficiency, less constrained selection on the UTR regions may have allowed the evolution of dynamic, stress-responsive structural elements. Protein binding to certain sequences also likely plays a role, and the interplay of RNA structure and protein un/binding will be an important avenue for future study.

We extended our analysis by developing a new concept, that of concordancy between transcript regions. The majority of mRNAs (~80%) are discordant, defined as two of the three different pairwise transcript regions changing in different directions under salt stress (Figure 6, Table 2C, S.7.c). If we consider that each coding sequence is flanked by two structural switches (the UTRs), this opens the possibility of three-level combinatorial regulation. However, it is notable that when all three regions (5’UTR, CDS, 3’UTR) gain or lose DMS reactivity in the same direction, which we define as ‘concordancy’, the effect of reactivity change on transcript abundance change is considerably more pronounced (Figure 4 vs. 6; Table 2A vs. 2C, S.6.a vs. S.7.c). This result suggests concerted protection from or exposure to various mechanisms of mRNA abundance regulation. Moreover, concordant change of reactivity between transcript regions is synergistic: when all transcript regions change in concert, the change of reactivity of the transcript as a whole (Figure 6 vs. Figure 4, leftmost panels) and as individual regions (Figure 6 vs. Figure 4; other panels) are more strongly correlated with the transcript’s change in abundance.

**Table 2.** Δ Mean Reactivity vs. Δ Abundance Correlations. Table 3A summarizes all the Δ mean reactivity vs. Δ abundance correlations, where 3B breaks it down by tissue specific (Unique) vs tissue nonspecific (Shared) transcripts. Table 3C provides the same information for fully concordant transcripts.

| Sample A | Sample B | Region | n | r | p |
| --- | --- | --- | --- | --- | --- |
| Shoot Control | Shoot Salt | whole | 15,875 | -0.73 | $< 2.2e^{-16}$ |
| | | 5'UTR | 14,435 | -0.29 | $< 2.2e^{-16}$ |
| | | CDS | 15,719 | -0.68 | $< 2.2e^{-16}$ |
| | | 3'UTR | 14,393 | -0.18 | $< 2.2e^{-16}$ |
| Root Control | Root Salt | whole | 15,725 | -0.58 | $< 2.2e^{-16}$ |
| | | 5'UTR | 14,334 | -0.16 | $< 2.2e^{-16}$ |
| | | CDS | 15,570 | -0.55 | $< 2.2e^{-16}$ |
| | | 3'UTR | 14,213 | -0.10 | $< 2.2e^{-16}$ |

B) Shared vs. Tissue Specific Transcripts, $\Delta$ Reactivity vs. $\Delta$ Abundance
| Tissue | Region | r All | r Shared | r Unique | p all | p shared | p unique |
| --- | --- | --- | --- | --- | --- | --- | --- |
| Shoot | Whole | -0.73 | -0.71 | -0.79 | $< 2.2e^{-16}$ | $< 2.2e^{-16}$ | $< 2.2e^{-16}$ |
| | 5'UTR | -0.29 | -0.27 | -0.35 | $< 2.2e^{-16}$ | $< 2.2e^{-16}$ | $< 2.2e^{-16}$ |
| | CDS | -0.68 | -0.65 | -0.76 | $< 2.2e^{-16}$ | $< 2.2e^{-16}$ | $< 2.2e^{-16}$ |
| | 3'UTR | -0.18 | -0.16 | -0.22 | $< 2.2e^{-16}$ | $< 2.2e^{-16}$ | $< 2.2e^{-16}$ |
| Root | Whole | -0.58 | -0.54 | -0.66 | $< 2.2e^{-16}$ | $< 2.2e^{-16}$ | $< 2.2e^{-16}$ |
| | 5'UTR | -0.16 | -0.14 | -0.23 | $< 2.2e^{-16}$ | $< 2.2e^{-16}$ | $< 2.2e^{-16}$ |
| | CDS | -0.55 | -0.49 | -0.65 | $< 2.2e^{-16}$ | $< 2.2e^{-16}$ | $< 2.2e^{-16}$ |
| | 3'UTR | -0.10 | -0.08 | -0.16 | $< 2.2e^{-16}$ | $< 2.2e^{-16}$ | $2.11e^{-13}$ |

C) $\Delta$ Mean Reactivity vs. $\Delta$ Abundance by Region, Fully Concordant
| Sample A | Sample B | $\Delta$ Reactivity | n | r | p |
| --- | --- | --- | --- | --- | --- |
| Shoot Control | Shoot Salt | whole | 3,155 | -0.82 | $< 2.2e^{-16}$ |
| | | 5'UTR | 3,155 | -0.58 | $< 2.2e^{-16}$ |
| | | CDS | 3,155 | -0.81 | $< 2.2e^{-16}$ |
| | | 3'UTR | 3,155 | -0.55 | $< 2.2e^{-16}$ |
| Root Control | Root Salt | whole | 3,175 | -0.65 | $< 2.2e^{-16}$ |
| | | 5'UTR | 3,175 | -0.43 | $< 2.2e^{-16}$ |
| | | CDS | 3,175 | -0.63 | $< 2.2e^{-16}$ |
| | | 3'UTR | 3,175 | -0.41 | $< 2.2e^{-16}$ |

Concordant reactivity increases may provide concerted access to both 5’ and 3’ exonucleases, as well as single-strand-specific endonucleases. The strong effect of concordant reactivity increases in promoting abundance decreases is consistent with known communication between 5’UTRs and 3’UTRs during stress (Vicens et al., 2018); for example, it is well-established that shortening of the poly A tail promotes decapping and subsequent 5’ to 3’ exonucleolytic degradation (Gallie, 1991, Muhlrad et al., 1994; Yamashita et al., 2005). That the 5’ and 3’ ends of the RNA are interacting is supported by comparison of panels N and O in Figure S.11. Conversely, concordant reactivity decreases may accompany protective mechanisms. For example, in mammalian systems is it known that sequestration of transcripts in stress granules, which is thought to protect transcripts from degradation, is associated with overall transcript compaction (folding) (Khong and Parker, 2018), which our assay would detect as a concordant decrease in DMS reactivity. It is notable that concordancy is significant not only between the ends of the RNA (i.e. in the 5’ and 3’ ends), but also between adjacent regions of the RNA in the 5’UTR and CDS, and the CDS and 3’UTR; for example, compare panels D and E, and panels I and J in Figure S.11. These results suggest that interaction of adjacent regions may play an underappreciated role in controlling transcript abundance.

It is of interest that concordant reactivity change in response to salt stress is tissue specific, i.e. only a small fraction (~23%) of mRNAs are fully concordant in both tissues, and an even smaller (~12%) set show concordancy in the same direction in both tissues (see Results for specific numbers). These observations support structural plasticity between tissues in their stress responses. Our GO analysis (Figure 8) of fully concordant transcripts under salinity stress shows enrichment in both tissues of encoded functions related to stress and to abiotic stimuli, particular for those transcripts that gained abundance and lost reactivity under salt stress, suggesting that structural mechanisms promote increased net expression of suites of stress-related transcripts in the two tissues. Our GO analysis of fully concordant transcripts that gain reactivity and lose abundance under salt stress in shoot shows a particular enrichment in genes related to photosynthesis (Figure 8). This structure-related loss in transcript abundance mirrors, and potentially contributes to, the well-known depression of photosynthesis under salt stress (Stepien and Johnson, 2009; Takahashi and Murata, 2008). In short, concerted structural change appears to retool the transcriptome to coordinate the physiological response to stress.

### mRNAs Exhibiting Dynamic Structures in Response to Salt are Enriched in Stress-Related Functions

In both tissues, transcripts with the largest changes of reactivity between control and salt stress were enriched in salt-stress, response to osmotic stress, ion binding, and post-translational protein modification gene ontology (GO) categories (Figure 7). These results indicate that much of the structural dynamism between conditions centers on genes that respond to stresses.

Using DMS reactivities as restraints in folding (Deigan et al. 2009; Ding et al., 2014; Su et al., 2018), we generated structures of 15,708 mRNAs in shoot and 15,557 mRNAs in root in both control and salt conditions. We used PPV to quantify the structural differences in these mRNAs between environmental conditions. The PPV distribution shows that transcripts on average had lower PPV values in root than in shoot (Figure 8A), suggesting that salt stress had a greater impact on RNA structure in roots than in shoots.

GO enrichment analysis of mRNAs in the bottom 5% of the PPV distribution, i.e. with large salt-induced changes in predicted structure, reveals over-representation of response to stimulus (GO 0050896) and protein transport (GO 0015031). The former category includes genes that function in salt stress or osmotic stress response, such as the illustrated rapid alkalinization factor 1 (RALF1), low temperature and salt responsive protein family genes (Figure 8B, 8E, 8F). Changes in response to stimulus may affect the function of stress responsive genes including RALF1, a peptide ligand that is perceived by the receptor-like kinase, FERONIA, with consequent modulation of the salinity response (Feng et al., 2018, Yu and Assmann, 2018). The latter category includes several Rab GTPases. Rab GTPase function in endosomal trafficking, and several have been implicated in plant salinity tolerance (Asaoka et al., 2013; Ebine et al., 2011; Yin et al., 2017). Given previous indications that RNA structure may affect protein folding during translation (Tang et al., 2016), salt-induced changes in RNA structure might have consequences for Rab functionality. In both tissues, transcripts in the top 5% of the PPV distribution, indicating predicted structural stability under salt stress, were enriched in mRNAs encoding proteins related to translation (P = 8.3e^−11^) and ribosome biogenesis (P = 8.12e^−13^) (Figure 8G,8H).

In a previous study (Ding et al., 2014), we compared in silico vs. in vivo structures of ~10,000 Arabidopsis seedling mRNAs, and found that those with low PPV, suggesting propensity to refold, were enriched in signaling and stress-related genes, while those with high PPV, suggesting structural stability, were enriched in housekeeping genes. That observation led to the hypotheses that signaling and stress-related genes are highly plastic in their structures as an intrinsic component of their environmental responsiveness, while housekeeping genes exhibit more stable structures, perhaps having evolved to maintain homeostasis under a wide range of conditions (Ding et al., 2014). Our present analyses, in which we directly compare *in vivo* structuromes of stressed vs. unstressed tissues, support those hypotheses. In the future, it would be interesting to evaluate shoot and root structuromes with a larger panel of stresses to evaluate the pervasiveness of these trends. Structurome data obtained under additional stress conditions could also reveal if some transcripts or transcript regions are consistently more prone to malleability between conditions while others are more static, which would suggest that the structurome has both fixed and dynamic features. Moreover, the biophysical nature of the changes remains to be explored; for instance, whether certain structural motifs are more prone to stress-specific refolding events or protein binding.

In summary, changes in transcript abundance, both within and between transcriptomes, appear intimately tied to changes in RNA structure. In the future, application of recent advances in our methodology, such as parallel probing with DMS and EDC, which is U-and G-specific (Wang et al. 2018; Mitchell et al., 2019), to report on absence of base-pairing for all four nucleobases, will allow an improved description of the dynamic RNA structurome. Such experiments may elucidate specific structure-related motifs, facilitating delineation of more specific patterns and molecular mechanisms that underlie structurome and transcriptome (Walley and Dehesh, 2010) reshaping by abiotic stresses. A deeper understanding of these mechanisms could augment efforts to engineer more salt-stress resistant crops (Cabello et al., 2014), and some structural elements have already been implicated as targets (Vashisht and Tuteja, 2006). Land mismanagement, use of marginal lands, and irrigation practices exacerbate salt stress and other pressures on crops, highlighting a need for fundamental understanding of how plants survive and thrive in challenging environments.

## Methods

### Plant growth and salt treatment

Arabidopsis accession Col-0 was used in this study. Sterilized Arabidopsis seeds were spread on agar plates containing 1⁄2 strength Murashige and Skoog (MS) medium with 1% sucrose and 0.8% agar (A1296; Sigma, St. Louis, MO, USA) and stratified at 4 °C in the dark for 2 days. After stratification, the plates were transferred to a growth chamber set at 21 °C with an 8 h light/16 h dark photoperiod with light intensity of 150 mol m^−2^ s^−1^and held vertically for 12 d. Seedlings were then transferred to a hydroponic setup as described in Yu and Assmann (2015) and grown hydroponically for another 12 days. The hydroponic growth solution was 1⁄4 strength Hoagland’s solution (0.25 mM KH_2_PO4, 3.71 μM FeNa-EDTA, 0.5 mM MgSO_4_, 1.26 mM KNO_3_, 1.26 mM Ca(NO_3_)_2_, 11.56 μM H_3_BO_3_, 2.29 μM MnCl_2_, 0.20 μM ZnCl_2_, 0.073 μM CuCl_2_, 0.026 μM Na_2_MoO_4_). Solutions were changed twice per week. After a total of 24 days of growth, half of the plants were exposed to salt stress by addition of 100 mM NaCl (final concentration) to the hydroponic solution, while the rest of the plants were maintained in the original solution. After 48 h of NaCl or control treatment, the plants were treated with DMS as described in a subsequent section below.

### Proline and ion content measurements

Twenty-four-day-old hydroponically grown seedlings were treated with or without 100 mM NaCl as above for 48 h. Shoots and roots were separated, gently blotted dry using paper towels, frozen in liquid nitrogen and dried in a CentriVap centrifugal vacuum concentrator. Dry weight of each sample was measured, and proline content was determined with a colorimetric assay (Ábrahám et al., 2010). Briefly, plant tissues were ground in 3% sulfosalicylic acid and centrifuged at 13,200 rpm for 5 min at room temperature using a benchtop centrifuge. Supernatant (100 μL) was transferred to a pre-made solution of 100 μL of 3% sulfosalicylic acid, 200 μL glacial acetic acid and 200 μL acidic ninhydrin. The reaction mixture was incubated at 96°C for 60 min, followed by addition of 1 mL toluene and vortexing to terminate the reaction. After centrifugation for 2 min at 10,000g, the absorbance at 520 nm of the chromophore in the toluene phase was measured with a spectrometer. The proline concentration was determined using a standard concentration curve of L-proline (Sigma) and calculated on a dry weight basis. Three independent replicates were performed with three biological samples in each replicate. Dry weight measurements are found in Table S.6. We also measured the levels of several ions on the same batch of seedlings as used for proline measurement (Table S.6.b) with three independent replicates, using inductively coupled plasma atomic emission spectroscopy as described previously (Yu and Assmann, 2015).

### *In vivo* DMS probing of *Arabidopsis* root and shoot tissue

To obtain “single-hit” modification kinetics (Su et al., 2018; Ding et al., 2014) in our DMS treatments, the duration of *in vivo* DMS treatment was first optimized. All DMS treatment procedures and RNA extraction procedures were conducted with proper safety equipment following the instructions in our publications (Ritchey et al. 2017; Su et al. 2018; Ritchey et al. 2019). For single-hit kinetics determination, we separately tested shoot and root tissue from Arabidopsis plants treated with/without 100 mM NaCl (control and salt stress). For the shoot tissue test, the shoots from five individual plants were used for each sample. For each condition, non-DMS-treated (-DMS) and DMS-treated (+DMS) samples were prepared. For the +DMS sample, the excised shoots were immersed in 20 mL DMS reaction buffer (40 mM HEPES (pH 7.5), 100 mM KCl, and 0.5 mM MgCl_2_) in a 50 mL conical centrifuge tube. Then 150 μl DMS (D186309, Sigma-Aldrich) was immediately added to the solution to a final concentration of 0.75% (~75 mM) with the following durations of DMS treatment: 1 min, 5 min, 10 min, 15 min and 20 min. After DMS treatment, dithiothreitol (DTT) at a final concentration of 0.5 M was supplied to quench DMS in the reaction for 2 min followed by two washes with deionized water before immediately drying and freezing the tissue in liquid nitrogen. The -DMS sample was processed through the same procedure by placing materials in the DMS buffer for 20 min but without addition of DMS. All procedures for the root tissue test were identical as for shoot treatments, except that the initial material was the root tissue from five plants, due to the lesser per plant mass of roots vs. shoots. RNA was extracted from frozen samples using the NucleoSpin RNA Plant kit (Cat# 740949, Macherey-Nagel, Germany) following the manufacturer’s protocol.

### Gene-specific reverse transcription for single-hit kinetics determination

Gene-specific reverse transcription (RT) of 18S rRNA was performed following the protocol of Ding et al. 2014. For each RT reaction, 1 μg total RNA in 5.5 μL total volume of RNase-free water was prepared. After adding 1 μL of ^32^P-radiolabeled primer (~250,000 cpm/μL) to target 18S rRNA (5′-AACTGATTTAATGAGCCATTCGCAG-3′), the solution was incubated at 75°C for 3 min, then cooled to 35°C. Then 2 μL of reverse transcription reaction buffer (5x) was added to a final concentration of 20 mM Tris–HCl (pH 7.5), 1 mM DTT, 100 mM KCl, 8 mM MgCl_2_, and 1 μL of 1 mM dNTPs. Annealing was allowed to proceed at 35°C for 5 min, after which the reaction solution was heated to 55°C for 1 min, 0.5 μL of Superscript III reverse transcriptase (Invitrogen; 100 U total) was added to the reaction, and the RT reaction was allowed to proceed at 55°C for 15 min. Next, 1 μL of 1 N NaOH was added to the solution, which was heated to 95 °C for 5 min to hydrolyze all RNAs and denature the reverse transcriptase. The RT product mixed with an equal volume of 2x loading dye was then loaded onto a 10% denaturing polyacrylamide gel (8.3 M urea) and run at 90 W for ~2 h. The gel was dried and exposed using a PhosphorImager (Molecular Dynamics) cassette. Gel images were collected with a Typhoon PhosphorImager 9410, and bands were quantified using ImageQuant 5.2. Based on these results, we chose 5 minutes DMS treatment for shoot and 1 minute DMS treatment for root to achieve similar single hit kinetics for the two distinct tissue types (Figure S.11). We then utilized these conditions for DMS treatment and subsequent Structure-seq library construction.

### Preparation of Structure-seq Libraries

After determination of single-hit kinetics for each condition, the plant materials for Structure-seq library generation were prepared. For each individual library, ~40 hydroponically grown Arabidopsis plants were used, which were dissected into shoot and root tissues. For the +DMS shoot sample, 40 excised shoots from control or NaCl treatment were immersed in 20 mL DMS reaction buffer (40 mM HEPES (pH 7.5), 100 mM KCl, and 0.5 mM MgCl_2_) in a 50 mL conical centrifuge tube. Then 150 μl DMS (D186309, Sigma-Aldrich) was immediately added to the solution to a final concentration of 0.75% (~75 mM) for 5 min of DMS treatment, after which dithiothreitol (DTT) at a final concentration of 0.5 M was supplied to quench DMS in the reaction for 2 min followed by two washes with deionized water before immediately drying and freezing the tissue in liquid nitrogen. The -DMS sample was processed through the same procedure by placing materials in the DMS buffer for 20 min but without addition of DMS. Three independent biological replicates were prepared for both control and NaCl-treated shoot libraries, and for both -DMS and +DMS libraries, for a total of three control -DMS libraries, three control +DMS libraries, three +NaCl −DMS libraries, and three +NaCl +DMS libraries. All procedures for generation of root Structure-seq libraries were identical as for shoot library preparation, starting with the root tissue from 40 plants each of control or NaCl treatment, except for a different duration of DMS treatment to maintain comparable single hit kinetics (Figure S.14). Frozen samples were subjected to RNA extraction using the NucleoSpin RNA Plant kit (Cat# 740949, Macherey-Nagel, Germany) following the manufacturer’s protocol.

Structure-seq libraries then were prepared according to the gel-based Structure-seq2 protocol (Ritchey et al. 2017; Su et al., 2018), and starting with ~250 μg total RNA. In total, 24 Structure-seq libraries were generated. Library cDNA size distribution and consistency between biological replicates were assessed from Bioanalyzer traces (Agilent 2100, Agilent Technologies). After qPCR to quantify library molarity, a pool of all libraries at equal molarity was made, and libraries were subjected to next-generation sequencing on an Illumina HiSeq 2500 at the Genomics Core Facility of the Penn State University to generate 150 nt single-end reads. Five HiSeq runs were performed to collect sufficient reads for downstream analyses, yielding a total of 1,109,349,671 reads. These libraries are available at the NCBI Gene Expression Omnibus (GEO) database as accession GSE124866.

### Data Handling and Processing

All sequenced libraries (FASTQ format) were first trimmed with Cutadapt (version 1.14) (Martin, 2011) via StructureFold2 (Tack et al., 2018) to remove 5’ and 3’ adapters, as well as low quality base calls on the 3’ ends of reads (libraries and trimming summarized in Table S.1). All individual sequencing runs of each library were combined (summarized in Table W.1) into a single FASTQ file representing a biological replicate before aligning to the TAIR10 (The Arabidopsis Information Resource) cDNA reference sans ribosomal sequences, using Bowtie2 (version 2.3.2) (Langmead and Salzberg, 2012) via StructureFold2 (Tack et al., 2018) (mappings summarized in Table W.2). These mappings were then post-processed using SAMtools to remove reads aligned to the opposite strand (version 0.1.19) (Li et al., 2009), as well as reads with more than 3 mismatches or indels compared to the reference, as well as any read with a mismatch at the first (5’) position of the read (Tack et al., 2018) (filtering summarized in Table W.3). Every filtered SAM file was then changed into an RTSC (Reverse Transcriptase Stop Count) file via Structurefold2 (Tack et al., 2018) before continuing with analysis. RT stop specificity was calculated from these RTSC files (Table W.4)

### DMS Library RT Stop Correlation Analyses

We pooled the RTSCs for all +DMS biological replicates of each condition to calculate total transcript coverage within that condition (Table W.1 for a description of all libraries) via Structurefold2 (Tack et al., 2018). We define coverage as the number of sequenced RT stops per A and C present in each transcript in the pool of all +DMS biological replicates of a condition. Thus, our coverage threshold of ≥ 1 mandates that we sequenced an average of one or more RT stop for each A or C residue of a given transcript within a condition in order to consider the structural information for that transcript to be resolvable (Kertesz et al., 2010). For all transcripts with coverage ≥ 1 in a condition, we correlated the component biological replicates within the condition in two ways. First, we performed all pairwise correlations between pairs of biological replicates on the entire transcriptome for each condition, i.e. correlated the number of RT stops on each A and C on every base in the transcriptome from transcripts above the combined coverage threshold for that condition using R (R Development Core Team, 2008) (cor.test) and plotting with ggplot (Wickham, 2009) (Figure S.2, hexbin plots). Second, we used the plyr (Wickham, 2011) package to organize our data such that we could perform a sub-correlation (cor.test()) for the RT stop values of each individual transcript between each biological replicate in each condition, plotting the distribution of each of these r values using ggplot (Wickham, 2009) (Figure S.2, violin plots). Taken together, these results indicate good replication between the individual biological replicates that make up our pools of data for each condition.

### Reactivity Calculations and Statistical Analyses

We calculated per base reactivity separately for each sample used in each of our contrasts using StructureFold2 (Tack et al., 2018), then subtracted pooled -DMS reads indicative of natural stops and natural stop-inducing modifications from corresponding +DMS modification RT stops (See StructureFold2 manual for equations, “rtsc to react module”). Every sample within each contrast was normalized to the same 2-8% normalization scale (Low and Weeks, 2010), where each scale contains separate 2-8% values for each transcript. The particular 2-8% scale used for normalization is relative as long as all data sets being contrasted are normalized to the same scale. This normalization of raw reactivity values from multiple samples by such a common scale (average reactivity of the top 8% of bases’ reactivities after ignoring the top 2%) ensures that all reactivity measurements between samples are relative to the same set of highly reactive bases. In both intra-tissue comparisons, the respective control scale was applied to the salt data, in both inter-tissue comparisons, the respective shoot scale was applied to the root data. The shoot control scale was used when comparing all four data sets. When there was no direct comparison between structuromes, e.g. when comparing abundance to mean reactivity (Figure S.6), each structurome’s internal 2-8% scale was used for normalization.

Transcript reactivities were summarized by the reactivity statistics module in StructureFold2, filtering the data to include only those transcripts with an overlapping coverage ≥ 1 in all samples present in a given contrast. As an additional filter, we restricted analysis to transcripts containing 10 or more As plus Cs before the last 30 bp of the transcript, such that mean reactivity values had a suitable number of measurements. These processed data can be found in supplemental item D; shoot control vs. salt (Tables D.1-D.4), root control vs. salt (Tables D.5-D8), control shoot vs. root (Tables D.9-D.12), salt shoot vs. root (Tables D.13-D.16), and four-way (Tables D.17-D.20) comparisons. We estimated transcript abundance via the StructureFold2 (Tack et al., 2018) rtsc abundance module, calculating TPM (Transcripts per kilobase-Million) from the reads in our corresponding pooled -DMS biological replicates for each sample. These data can be found in Tables D.21-24. Unless specified, all downstream analysis was done with R (R Development Core Team, 2008) and plots were generated via the ggplot (Wickham, 2009) package. Base R functionality was used for all tests and analyses; all t-tests were carried out using t.test(), all correlations carried out using cor.test(), aov() was used for ANOVA and information was extracted with summary() and TukeyHSD(). StructureFold2 formats all such data to be readily used by R. Concordant transcripts and regions were grouped by comparing the reactivity change of transcript regions between conditions via subset(), i.e. regions that shared the direction of reactivity change were considered concordant.

### GO Analyses

To run GO analysis on transcripts with large reactivity change between conditions or tissues, transcript subsets were quantitatively selected from our data, collecting the 5% top and bottom extremes of mean reactivity change between conditions or tissues. For the GO analyses on concordant transcripts, we qualitatively subset our transcripts into those that increased in abundance and lost reactivity (concordant protection) and those that decreased in abundance and increased in reactivity (concordant exposure). For the GO analyses using PPV, we selected the 5% top and bottom extremes of PPV between conditions. All transcript subsets were then reduced to reflect their gene locus of origin before using these to run GO analyses via agriGOv2 (Tian et al., 2017). These individual results were then combined via a custom Python script back into a convenient csv for easy analysis and plotting of several GO analyses simultaneously using R (R Development Core Team, 2008) and ggplot (Wickham, 2009). Altogether, we ran 32 GO analyses involving reactivity change (Tables G.1-G.32), 4 involving concordancy (Tables G.33-G.36) and 4 involving PPV (Tables G.37-40), for a total of 40 analyses.

### ANOVAs of Mean Reactivity

To contrast all of our structuromes together, we implemented a series of large scale ANOVAs. We used the 2-8% normalization scale from shoot control and also applied it to shoot salt, root control, and root salt as a common normalization scale such that all the data would be comparable, and used only transcripts that had a coverage ≥ 1 in the corresponding +DMS sample of every condition (Figure 1F, Figure S.5). This allowed four two-way ANOVAs (whole transcript: n=12,978, 5’UTR: n=12,117 UTRs, CDS: n=12,870, 3’UTR: n=12,278) comparing the mean reactivity of whole transcript and each region against their treatment and tissue (mean reactivity ~ treatment * tissue). We combined the separate transcript regions, and ran a three-way ANOVA such that region could also be used as determinant of mean reactivity (mean reactivity ~ treatment * tissue * region) (Figure S.5, Table S.5). Tissue and treatment:tissue were always stronger effects than treatment alone. When ‘region’ was included as a factor, it was also a stronger determinant than treatment.

### Structure Prediction and Derived Statistics

Reactivity files generated from intra-tissue comparisons were used as restraints to guide *in silico* folding of each transcript and transcript region above the coverage threshold in both control and NaCl conditions within a tissue via the RNAStructure Fold program (Reuter and Matthews, 2010) with StructureFold2 (Tack et al., 2018). The output is a predicted minimum free energy structure for each transcript and transcript region. We then calculated PPV (Positive Predictive Value) between each pair of in vivo restrained folds by driving the RNAStructure scorer program with StructureFold2, thereby assaying how similar the folds of each transcript/region were between conditions (Tables F.1-F.8). Circle plots between folds were generated with CircleCompare (Reuter and Matthews, 2010) and GO analysis was performed on the top and bottom 5% most extreme PPV values in both tissues with AgriGOv2 (Tian et al., 2017).

### Comparison to Anderson et al. data

We extracted the salt-stress induced transcript fold changes from the data supplement (S3) of Anderson et al. (2018). These fold changes were used as overlay colors on our delta mean reactivity versus delta transcript abundance plots (Figure 10), showing high parity of salinity-induced abundance changes between Anderson et al. (2018) and our study.

## Supporting information

Supplemental Workflow (W)

Supplemental Raw (R)

Supplemental Folding (F)

## Author Contributions and Acknowledgements

S.M.A. and P.C.B. conceived the study. Z.S. performed the experiments, excluding measurements of ion concentrations and proline, which were performed by Y.Y.. D.C.T. performed all structurome and statistical analyses. D.C.T. and Z.S. performed GO analyses. S.M.A., D.C.T., P.C.B., and Z.S. contributed ideas, discussed the results, and wrote the manuscript. All authors edited the manuscript. We thank Dr. Elizabeth Jolley and Megan Sylvia for reading over the manuscript and giving feedback. This project was supported by NSF grant IOS-1339282 to P.C.B. and S.M.A..

## References

Abogadallah, G. M. (2010). Insights into the significance of antioxidative defense under salt stress. Plant signaling & behavior, 5(4), 369–374.

Ábrahám, E., Hourton-Cabassa, C., Erdei, L., & Szabados, L. (2010). Methods for determination of proline in plants. In Plant Stress Tolerance (pp. 317–331). Humana Press.

Anderson, S. J., Kramer, M. C., Gosai, S. J., Yu, X., Vandivier, L. E., Nelson, A. D., … & Gregory, B. D. (2018). N6-methyladenosine inhibits local ribonucleolytic cleavage to stabilize mRNAs in Arabidopsis. Cell reports, 25(5), 1146–1157.

Asaoka, R., Uemura, T., Ito, J., Fujimoto, M., Ito, E., Ueda, T., & Nakano, A. (2013). Arabidopsis RABA1 GTPases are involved in transport between the trans-Golgi network and the plasma membrane, and are required for salinity stress tolerance. The Plant Journal, 73(2), 240–249.

Apse, M. P., & Blumwald, E. (2007). Na*^+^* transport in plants. FEBS letters, 581(12), 2247–2254.

Bahieldin, A., Atef, A., Sabir, J. S., Gadalla, N. O., Edris, S., Alzohairy, A. M., … & Hassan, S. M. (2015). RNA-Seq analysis of the wild barley (H. spontaneum) leaf transcriptome under salt stress. Comptes rendus biologies, 338(5), 285–297.

Bevilacqua, P. C., Ritchey, L. E., Su, Z., & Assmann, S. M. (2016). Genome-wide analysis of RNA secondary structure. Annual review of genetics, 50, 235–266.

Bloomfield, V. A., Crothers, D. M. & Tinoco, I., Jr. (2000). Nucleic Acids: Structures, Properties, and Functions, University Science Books, Sausalito, California.

Carey, J. N., Mettert, E. L., Roggiani, M., Myers, K. S., Kiley, P. J., & Goulian, M. (2018). Regulated Stochasticity in a Bacterial Signaling Network Permits Tolerance to a Rapid Environmental Change. Cell, 173(1), 196–207.

Cabello, J. V., Lodeyro, A. F., & Zurbriggen, M. D. (2014). Novel perspectives for the engineering of abiotic stress tolerance in plants. Current Opinion in Biotechnology, 26, 62–70.

Cui, J., Ren, G., Qiao, H., Xiang, X., Huang, L., & Chang, J. (2018). Comparative Transcriptome Analysis of Seedling Stage of Two Sorghum Cultivars Under Salt Stress. Journal of plant growth regulation, 37(3), 986–998.

Culotta, V. C., Yang, M., & O’Halloran, T. V. (2006). Activation of superoxide dismutases: putting the metal to the pedal. Biochimica et Biophysica Acta (BBA)-Molecular Cell Research, 1763(7), 747–758.

Deigan, K. E., Li, T. W., Mathews, D. H., & Weeks, K. M. (2009). Accurate SHAPE-directed RNA structure determination. Proceedings of the National Academy of Sciences, 106(1), 97–102.

Deinlein, U., Stephan, A. B., Horie, T., Luo, W., Xu, G., & Schroeder, J. I. (2014). Plant salt-tolerance mechanisms. Trends in plant science, 19(6), 371–379.

Deng, H., Cheema, J., Zhang, H., Woolfenden, H., Norris, M., Liu, Z., … & Cao, X. (2018). Rice in vivo RNA structurome reveals RNA secondary structure conservation and divergence in plants. Molecular plant, 11(4), 607–622.

Ding, Y., Tang, Y., Kwok, C. K., Zhang, Y., Bevilacqua, P. C., & Assmann, S. M. (2014). In vivo genome-wide profiling of RNA secondary structure reveals novel regulatory features. Nature, 505(7485), 696.

Ding, W., Fang, W., Shi, S., Zhao, Y., Li, X., & Xiao, K. (2016). Wheat WRKY type transcription factor gene TaWRKY1 is essential in mediating drought tolerance associated with an ABA-dependent pathway. Plant molecular biology reporter, 34(6), 1111–1126.

Dinneny, J. R., Long, T. A., Wang, J. Y., Jung, J. W., Mace, D., Pointer, S., … & Benfey, P. N. (2008). Cell identity mediates the response of Arabidopsis roots to abiotic stress. Science, 320(5878), 942–945.

Ebine, K., Fujimoto, M., Okatani, Y., Nishiyama, T., Goh, T., Ito, E., … & Thordal-Christensen, H. (2011). A membrane trafficking pathway regulated by the plant-specific RAB GTPase ARA6. Nature Cell Biology, 13(7), 853.

Feng, J., Li, J., Gao, Z., Lu, Y., Yu, J., Zheng, Q., … & Zhu, Z. (2015). SKIP confers osmotic tolerance during salt stress by controlling alternative gene splicing in Arabidopsis. Molecular plant, 8(7), 1038–1052.

Feng, W., Kita, D., Peaucelle, A., Cartwright, H. N., Doan, V., Duan, Q., … & Yvon, R. (2018). The FERONIA receptor kinase maintains cell-wall integrity during salt stress through Ca2+ signaling. Current Biology, 28(5), 666–675.

Floris, M., Mahgoub, H., Lanet, E., Robaglia, C., & Menand, B. (2009). Post-transcriptional regulation of gene expression in plants during abiotic stress. International journal of molecular sciences, 10(7), 3168–3185.

Food and Agriculture Organization of the United Nations (FAO). (2009). How to feed the world in 2050.

Gallie, D. R. (1991). The cap and poly (A) tail function synergistically to regulate mRNA translational efficiency. Genes & development, 5(11), 2108–2116.

Gu, J., Xia, Z., Luo, Y., Jiang, X., Qian, B., Xie, H., … & Wang, Z. Y. (2017). Spliceosomal protein U1A is involved in alternative splicing and salt stress tolerance in Arabidopsis thaliana. Nucleic acids research, 46(4), 1777–1792.

Heilman-Miller, S. L., Thirumalai, D., & Woodson, S. A. (2001). Role of counterion condensation in folding of the Tetrahymena ribozyme. I. Equilibrium stabilization by cations. Journal of molecular biology, 306(5), 1157–1166.

Huot, B., Yao, J., Montgomery, B. L., & He, S. Y. (2014). Growth–defense tradeoffs in plants: a balancing act to optimize fitness. Molecular plant, 7(8), 1267–1287.

Jiao, Y., Riechmann, J. L., & Meyerowitz, E. M. (2008). Transcriptome-wide analysis of uncapped mRNAs in Arabidopsis reveals regulation of mRNA degradation. The Plant Cell, 20(10), 2571–2585.

Jinek, M., Coyle, S. M., & Doudna, J. A. (2011). Coupled 5′ nucleotide recognition and processivity in Xrn1-mediated mRNA decay. Molecular cell, 41(5), 600–608.

Kawa, D., & Testerink, C. (2017). Regulation of mRNA decay in plant responses to salt and osmotic stress. Cellular and Molecular Life Sciences, 74(7), 1165–1176.

Kertesz, M., Wan, Y., Mazor, E., Rinn, J. L., Nutter, R. C., Chang, H. Y., & Segal, E. (2010). Genome-wide measurement of RNA secondary structure in yeast. Nature, 467(7311), 103.

Khong, A., & Parker, R. (2018). mRNP architecture in translating and stress conditions reveals an ordered pathway of mRNP compaction. J Cell Biol, 217(12), 4124–4140.

Kreps, J. A., Wu, Y., Chang, H. S., Zhu, T., Wang, X., & Harper, J. F. (2002). Transcriptome changes for Arabidopsis in response to salt, osmotic, and cold stress. Plant physiology, 130(4), 2129–2141.

Lambert, D., & Draper, D. E. (2007). Effects of osmolytes on RNA secondary and tertiary structure stabilities and RNA-Mg2+ interactions. Journal of molecular biology, 370(5), 993–1005.

Langmead, B., & Salzberg, S. L. (2012). Fast gapped-read alignment with Bowtie 2. Nature methods, 9(4), 357.

Leamy, K. A., Assmann, S. M., Mathews, D. H., & Bevilacqua, P. C. (2016). Bridging the gap between in vitro and in vivo RNA folding. Quarterly reviews of biophysics, 49.

Li, H., Handsaker, B., Wysoker, A., Fennell, T., Ruan, J., Homer, N., … & Durbin, R. (2009). The sequence alignment/map format and SAMtools. Bioinformatics, 25(16), 2078–2079.

Li, F., Zheng, Q., Vandivier, L. E., Willmann, M. R., Chen, Y., & Gregory, B. D. (2012). Regulatory impact of RNA secondary structure across the Arabidopsis transcriptome. The Plant Cell, 24(11), 4346–4359.

Liang, W., Ma, X., Wan, P., & Liu, L. (2018). Plant salt-tolerance mechanism: A review. Biochemical and biophysical research communications, 495(1), 286–291.

Lin-Chao, S., Chiou, N. T., & Schuster, G. (2007). The PNPase, exosome and RNA helicases as the building components of evolutionarily-conserved RNA degradation machines. Journal of biomedical science, 14(4), 523–532.

Low, J. T., & Weeks, K. M. (2010). SHAPE-directed RNA secondary structure prediction. Methods, 52(2), 150–158.

Mandal, M., & Breaker, R. R. (2004). Gene regulation by riboswitches. Nature reviews Molecular cell biology, 5(6), 451.

Manning, G. S. (1972). On the Application of Polyelectrolyte “Limiting Laws” to the Helix-Coil Transition of DNA. II. The Effect of Mg++ Counterions. Biopolymers: Original Research on Biomolecules, 11(5), 951–955.

Mattioli, R., Costantino, P., & Trovato, M. (2009). Proline accumulation in plants: not only stress. Plant signaling & behavior, 4(11), 1016–1018.

Martin, M. (2011). Cutadapt removes adapter sequences from high-throughput sequencing reads. EMBnet. journal, 17(1), pp-10.

Merret, R., Descombin, J., Juan, Y. T., Favory, J. J., Carpentier, M. C., Chaparro, C., … & Bousquet-Antonelli, C. (2013). XRN4 and LARP1 are required for a heat-triggered mRNA decay pathway involved in plant acclimation and survival during thermal stress. Cell reports, 5(5), 1279–1293.

Merret, R., Nagarajan, V. K., Carpentier, M. C., Park, S., Favory, J. J., Descombin, J., … & Bousquet-Antonelli, C. (2015). Heat-induced ribosome pausing triggers mRNA co-translational decay in Arabidopsis thaliana. Nucleic acids research, 43(8), 4121–4132.

Mitchell, D., Ritchey, L. E., Park, H., Babitzke, P., Assmann, S. M., & Bevilacqua, P. C. (2018). Glyoxals as in vivo RNA structural probes of guanine base-pairing. RNA, 24(1), 114–124.

Mitchell, D., Renda, A. J., Douds, C. A., Babitzke, P., Assmann, S. M., & Bevilacqua, P. C. (2019). In vivo RNA structural probing of uracil and guanine base-pairing by 1-ethyl-3-(3-dimethylaminopropyl) carbodiimide (EDC). RNA, 25(1), 147–157.

Molin, C., Jauhiainen, A., Warringer, J., Nerman, O., & Sunnerhagen, P. (2009). mRNA stability changes precede changes in steady-state mRNA amounts during hyperosmotic stress. Rna.

Muhlrad, D., Decker, C. J., & Parker, R. (1994). Deadenylation of the unstable mRNA encoded by the yeast MFA2 gene leads to decapping followed by 5’--> 3’digestion of the transcript. Genes & development, 8(7), 855–866.

Munns, R., & Tester, M. (2008). Mechanisms of salinity tolerance. Annu. Rev. Plant Biol., 59, 651–681.

Narsai, R., Howell, K. A., Millar, A. H., O’Toole, N., Small, I., & Whelan, J. (2007). Genome-wide analysis of mRNA decay rates and their determinants in Arabidopsis thaliana. The Plant Cell, 19(11), 3418–3436.

Narberhaus, F., Waldminghaus, T., & Chowdhury, S. (2006). RNA thermometers. FEMS microbiology reviews, 30(1), 3–16.

Nilsen, T. W., & Graveley, B. R. (2010). Expansion of the eukaryotic proteome by alternative splicing. Nature, 463(7280), 457.

Parker, R., & Sheth, U. (2007). P bodies and the control of mRNA translation and degradation. Molecular cell, 25(5), 635–646.

Pennisi, E. (2018). Development cell by cell. Science Magazine. (Link)

Qadir, M., Quillérou, E., Nangia, V., Murtaza, G., Singh, M., Thomas, R. J., … & Noble, A. D. (2014, November). Economics of salt-induced land degradation and restoration. In Natural Resources Forum (Vol. 38, No. 4, pp. 282–295).

R Development Core Team. R: A language and environment for statistical computation, 2008. http://www.R-project.org.

Rabani, M., Pieper, L., Chew, G. L., & Schier, A. F. (2017). A Massively Parallel Reporter Assay of 3′ UTR Sequences Identifies In Vivo Rules for mRNA Degradation. Molecular cell, 68(6), 1083–1094.

Rengasamy, P. (2006). World salinization with emphasis on Australia. Journal of experimental botany, 57(5), 1017–1023.

Reuter, J. S., & Mathews, D. H. (2010). RNAstructure: software for RNA secondary structure prediction and analysis. BMC bioinformatics, 11(1), 129.

Ritchey, L. E., Su, Z., Tang, Y., Tack, D. C., Assmann, S. M., & Bevilacqua, P. C. (2017). Structure-seq2: sensitive and accurate genome-wide profiling of RNA structure in vivo. Nucleic acids research, 45(14), e135–e135.

Ritchey LE, Su Z, Assmann SM, Bevilacqua PC (2019) In vivo genome-wide RNA structure probing with structure-seq. In: Chekanova JA, Wang H.-L. V. (eds) Plant long non-coding RNAs: methods and protocols, Methods in molecular biology, vol 1933. Springer, New York, pp 305–341

Romero-Santacreu, L., Moreno, J., Pérez-Ortín, J. E., & Alepuz, P. (2009). Specific and global regulation of mRNA stability during osmotic stress in Saccharomyces cerevisiae. Rna.

Rouskin, S., Zubradt, M., Washietl, S., Kellis, M., & Weissman, J. S. (2014). Genome-wide probing of RNA structure reveals active unfolding of mRNA structures in vivo. Nature, 505(7485), 701.

Roy, S. J., Negrão, S., & Tester, M. (2014). Salt resistant crop plants. Current Opinion in Biotechnology, 26, 115–124.

Shinozaki, K., Yamaguchi-Shinozaki, K., & Seki, M. (2003). Regulatory network of gene expression in the drought and cold stress responses. Current opinion in plant biology, 6(5), 410–417.

Stepien, P., & Johnson, G. N. (2009). Contrasting responses of photosynthesis to salt stress in the glycophyte Arabidopsis and the halophyte Thellungiella: role of the plastid terminal oxidase as an alternative electron sink. Plant physiology, 149(2), 1154–1165.

Sorenson, R. S., Deshotel, M. J., Johnson, K., Adler, F. R., & Sieburth, L. E. (2018). Arabidopsis mRNA decay landscape arises from specialized RNA decay substrates, decapping-mediated feedback, and redundancy. Proceedings of the National Academy of Sciences, 115(7), E1485–E1494.

Spitale, R. C., Flynn, R. A., Zhang, Q. C., Crisalli, P., Lee, B., Jung, J. W., … & Chang, H. Y. (2015). Structural imprints in vivo decode RNA regulatory mechanisms. Nature, 519(7544), 486.

Su, Z., Tang, Y., Ritchey, L. E., Tack, D. C., Zhu, M., Bevilacqua, P. C., & Assmann, S. M. (2018). Genome-wide RNA structurome reprogramming by acute heat shock globally regulates mRNA abundance. Proceedings of the National Academy of Sciences, 115(48), 12170–12175.

Sun, W., Xu, X., Zhu, H., Liu, A., Liu, L., Li, J., & Hua, X. (2010). Comparative transcriptomic profiling of a salt-tolerant wild tomato species and a salt-sensitive tomato cultivar. Plant and Cell Physiology, 51(6), 997–1006.

Sun, Y., Kong, X., Li, C., Liu, Y., & Ding, Z. (2015). Potassium retention under salt stress is associated with natural variation in salinity tolerance among Arabidopsis accessions. PLoS One, 10(5), e0124032.

Tack, D. C., Tang, Y., Ritchey, L. E., Assmann, S. M., & Bevilacqua, P. C. (2018). StructureFold2: Bringing chemical probing data into the computational fold of RNA structural analysis. Methods.

Tang, Y., Assmann, S. M., & Bevilacqua, P. C. (2016). Protein structure is related to RNA structural reactivity in vivo. Journal of molecular biology, 428(5), 758–766.

Takahashi, S., & Murata, N. (2008). How do environmental stresses accelerate photoinhibition?. Trends in plant science, 13(4), 178–182.

Tan, Z. J., & Chen, S. J. (2011). Salt contribution to RNA tertiary structure folding stability. Biophysical journal, 101(1), 176–187.

Teakle, N. L., & Tyerman, S. D. (2010). Mechanisms of Cl-transport contributing to salt tolerance. Plant, Cell & Environment, 33(4), 566–589.

Tester, M., & Davenport, R. (2003). Na+ tolerance and Na+ transport in higher plants. Annals of botany, 91(5), 503–527.

The Arabidopsis Information Resource (TAIR), https://www.arabidopsis.org/download/index-auto.jsp?dir=%2Fdownload_files%2FSequences%2FTAIR10_blastsets, on www.arabidopsis.org, [4/20/2016].

Tian, T., Liu, Y., Yan, H., You, Q., Yi, X., Du, Z., Xu, W., & Su, Z. (2017) agriGO v2.0: a GO analysis toolkit for the agricultural community, 2017 update. Nuc. Acids. Res. 45: W122–W129.

Tijerina, P., Mohr, S., & Russell, R. (2007). DMS footprinting of structured RNAs and RNA–protein complexes. Nature protocols, 2(10), 2608.

Trinchant, J. C., Boscari, A., Spennato, G., Van de Sype, G., & Le Rudulier, D. (2004). Proline betaine accumulation and metabolism in alfalfa plants under sodium chloride stress. Exploring its compartmentalization in nodules. Plant physiology, 135(3), 1583–1594.

Tomecki, R., Kristiansen, M. S., Lykke-Andersen, S., Chlebowski, A., Larsen, K. M., Szczesny, R. J., … & Dziembowski, A. (2010). The human core exosome interacts with differentially localized processive RNases: hDIS3 and hDIS3L. The EMBO journal, 29(14), 2342–2357

Vashisht, A. A., & Tuteja, N. (2006). Stress responsive DEAD-box helicases: a new pathway to engineer plant stress tolerance. Journal of Photochemistry and Photobiology B: Biology, 84(2), 150–160.

Vicens, Q., Kieft, J. S., & Rissland, O. S. (2018). Revisiting the Closed-Loop Model and the Nature of mRNA 5′–3′ Communication. Molecular cell, 72(5), 805–812.

Vu, D. T., Yamada, T., & Ishidaira, H. (2018). Assessing the impact of sea level rise due to climate change on seawater intrusion in Mekong Delta, Vietnam. Water Science and Technology, 77(6), 1632–1639.

Wachter, A., Tunc-Ozdemir, M., Grove, B. C., Green, P. J., Shintani, D. K., & Breaker, R. R. (2007). Riboswitch control of gene expression in plants by splicing and alternative 3′ end processing of mRNAs. The Plant Cell, 19(11), 3437–3450.

Walley, J. W., & Dehesh, K. (2010). Molecular mechanisms regulating rapid stress signaling networks in Arabidopsis. Journal of integrative plant biology, 52(4), 354–359.

Wan, Y., Qu, K., Ouyang, Z., Kertesz, M., Li, J., Tibshirani, R., … & Chang, H. Y. (2012). Genome-wide measurement of RNA folding energies. Molecular cell, 48(2), 169–181.

Wan, Y., Qu, K., Zhang, Q. C., Flynn, R. A., Manor, O., Ouyang, Z., … & Chang, H. Y. (2014). Landscape and variation of RNA secondary structure across the human transcriptome. Nature, 505(7485), 706.

Wang, P. Y., Sexton, A. N., Culligan, W. J., & Simon, M. D. (2019). Carbodiimide reagents for the chemical probing of RNA structure in cells. RNA, 25(1), 135–146.

Wang, J., Zhu, J., Zhang, Y., Fan, F., Li, W., Wang, F., … & Yang, J. (2018). Comparative transcriptome analysis reveals molecular response to salinity stress of salt-tolerant and sensitive genotypes of indica rice at seedling stage. Scientific reports, 8(1), 2085.

Wickham, H. (2011). The split-apply-combine strategy for data analysis. Journal of Statistical Software, 40(1), 1–29.

Wickham, H. (2016). ggplot2: elegant graphics for data analysis. Springer-Verlag New York, 2009.

Wilkinson, K. A., Merino, E. J., & Weeks, K. M. (2006). Selective 2′-hydroxyl acylation analyzed by primer extension (SHAPE): quantitative RNA structure analysis at single nucleotide resolution. Nature protocols, 1(3), 1610.

Yamashita, A., Chang, T. C., Yamashita, Y., Zhu, W., Zhong, Z., Chen, C. Y. A., & Shyu, A. B. (2005). Concerted action of poly (A) nucleases and decapping enzyme in mammalian mRNA turnover. Nature structural & molecular biology, 12(12), 1054.

Yin, C., Karim, S., Zhang, H., & Aronsson, H. (2017). Arabidopsis RabF1 (ARA6) is involved in salt stress and dark-induced senescence (DIS). International journal of molecular sciences, 18(2), 309.

Yu, Y., & Assmann, S. M. (2015). The heterotrimeric G-protein β subunit, AGB 1, plays multiple roles in the A rabidopsis salinity response. Plant, cell & environment, 38(10), 2143–2156.

Yu, Y., & Assmann, S. M. (2018). Inter-relationships between the heterotrimeric Gβ subunit AGB1, the receptor-like kinase FERONIA, and RALF1 in salinity response. Plant, cell & environment, 41(10), 2475–2489.

Zhang, M., Kong, X., Xu, X., Li, C., Tian, H., & Ding, Z. (2015). Comparative transcriptome profiling of the maize primary, crown and seminal root in response to salinity stress. PloS one, 10(3), e0121222.

Zimeri, A. M., Dhankher, O. P., McCaig, B., & Meagher, R. B. (2005). The plant MT1 metallothioneins are stabilized by binding cadmiums and are required for cadmium tolerance and accumulation. Plant molecular biology, 58(6), 839–855.

Zhu, G., Li, W., Zhang, F., & Guo, W. (2018). RNA-seq analysis reveals alternative splicing under salt stress in cotton, Gossypium davidsonii. BMC genomics, 19(1), 73.

Ziehler, W. A., & Engelke, D. R. (2000). Probing RNA structure with chemical reagents and enzymes. Current protocols in nucleic acid chemistry, (1), 6–1.

