## Supplemental Figures for "Tissue-specific changes in the RNA structurome mediate salinity response in *Arabidopsis*"

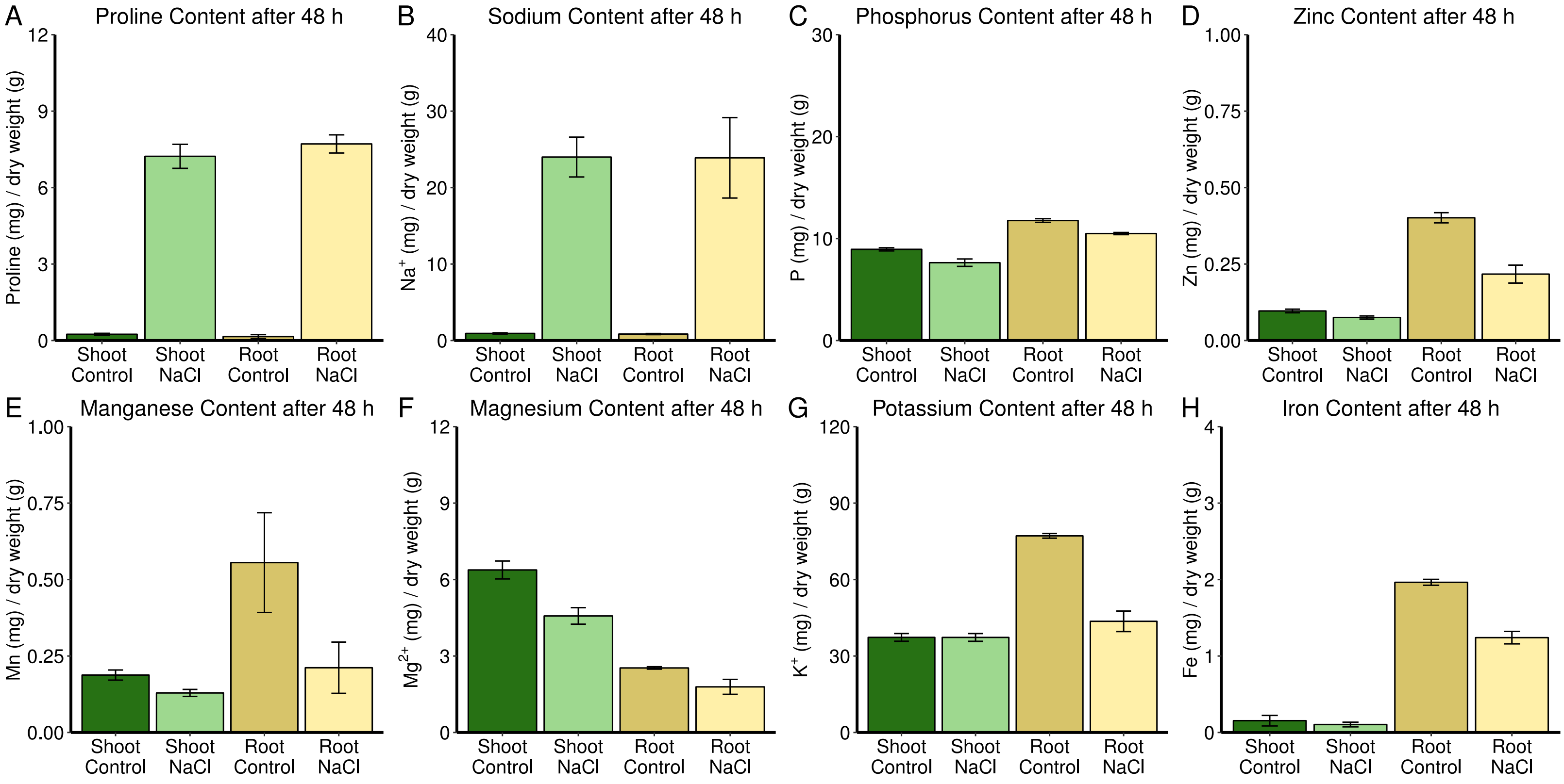


**Figure S.1 - The Effect of Salt Stress on Arabidopsis Ion Content**
Measurements of proline, sodium, phosphorus, zinc, manganese, magnesium, and iron content under control conditions and after 48 h of salt stress (100 mM NaCl) are summarized as bar graphs. Charges are not designated for phosphorus, zinc, manganese, and iron due to the potential for various valences to exist within the cell. All accompanying measurements and statistics can be found in Table S.1 and analysis via ANOVA can be found in Table S.2.

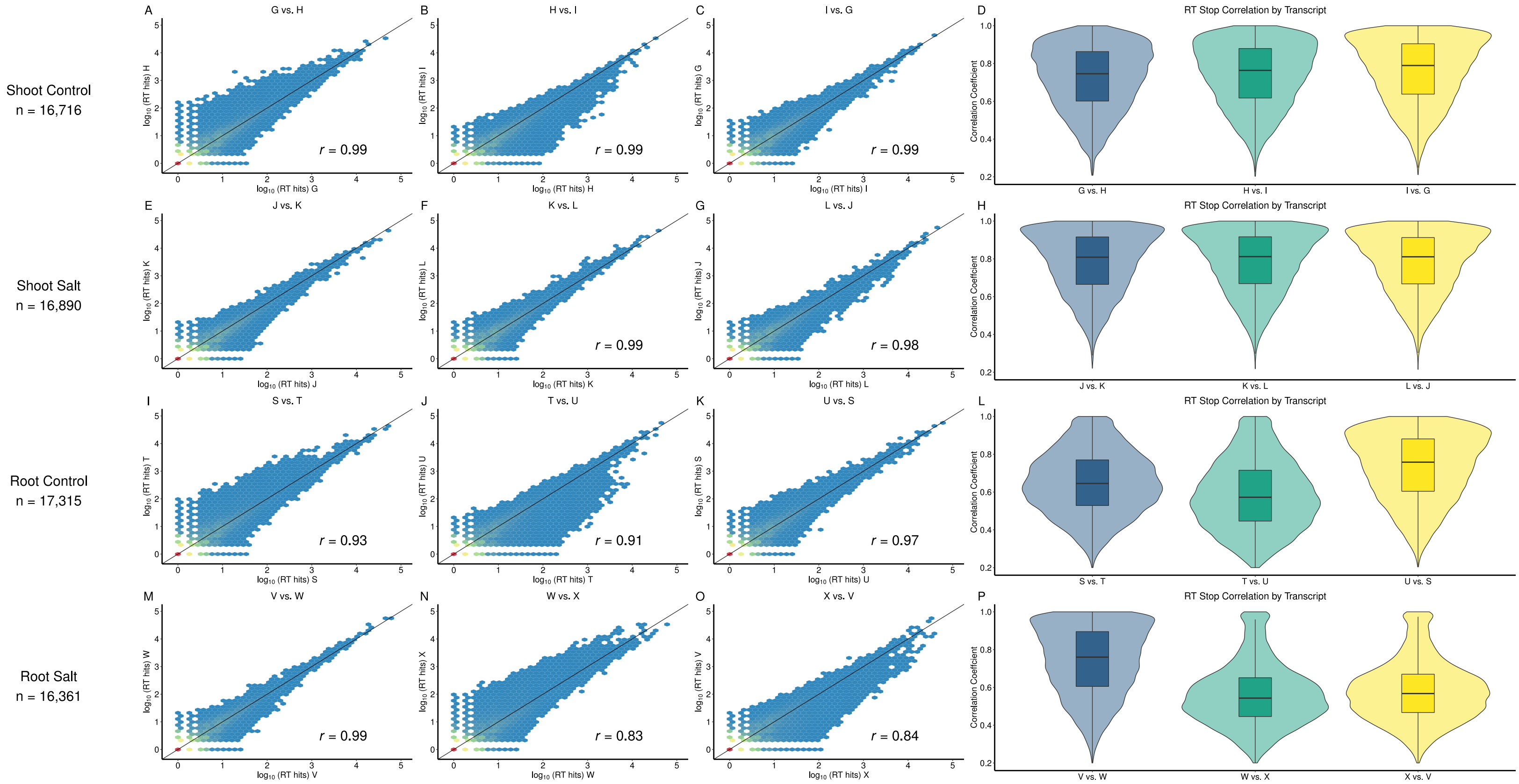


**Figure S.2 - Replicate Repeatability Correlations**

A-C,E-G,I-K,M-O) Whole transcriptome correlations of A/C nucleotide reverse transcriptase stop counts in DMS-treated libraries between biological replicates of the same condition: shoot contol, shoot salt, root control, and root salt respectively. D,H,L,P) Distributions of per transcript correlations of A,C nucleotide reverse transcriptase stop counts between biological replicates of the same condition. DMS modifications are suitably repeatable between our biological replicates on both a whole transcriptome and individual transcript basis to proceed with analysis. A complete table of all libraries and treatments is included in Table W.1. Numbers of transcripts used in this analysis are higher than those in Figure 1F due to some transcripts not having the requisite 10 or more As plus Cs before the last 30 bp of the transcript, i.e. correlation of stops could be assessed but not a reliable mean reactivity.

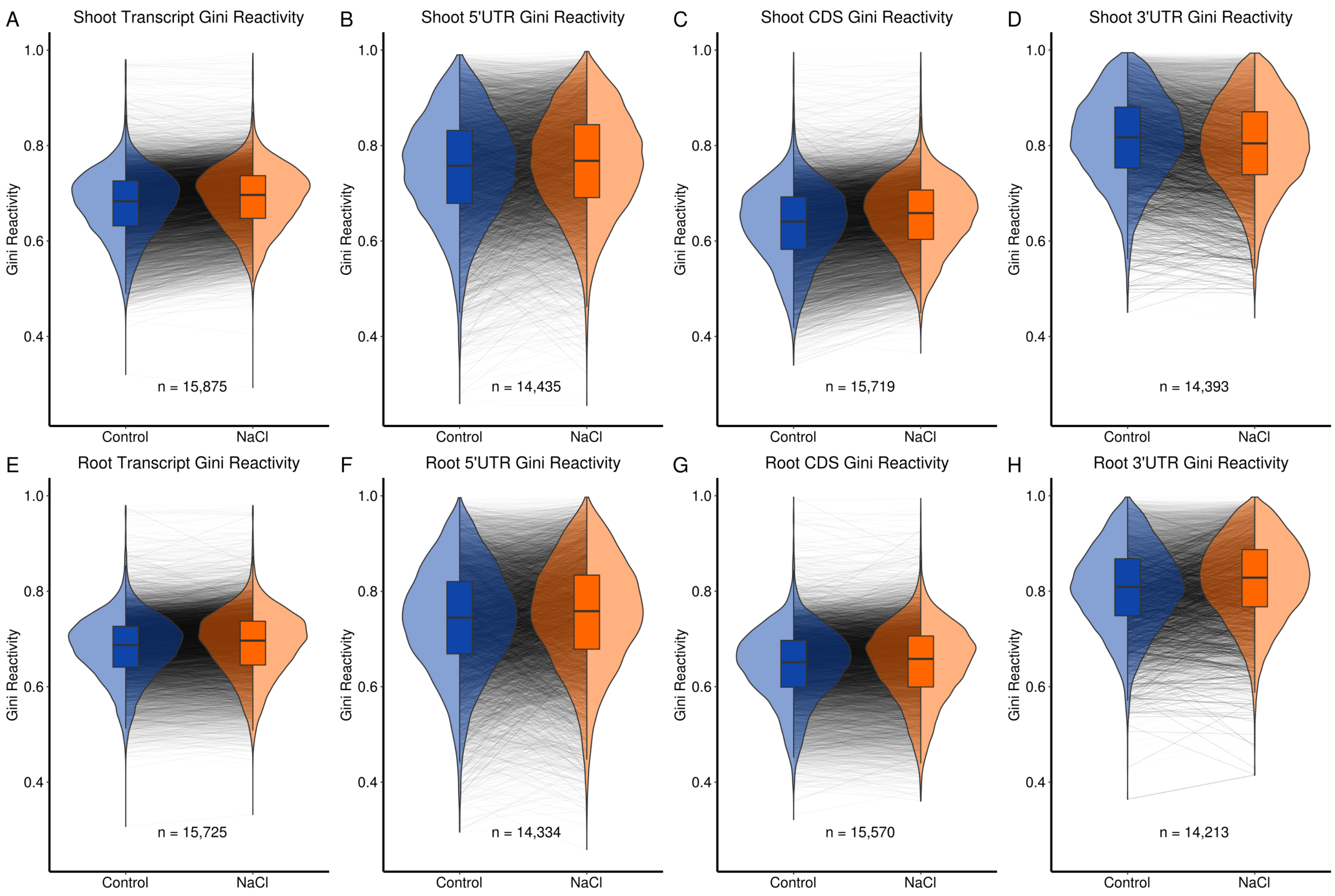


**Figure S.3 - The Effect of Salt Stress on Arabidopsis Intra-Tissue Gini Index of Transcript Reactivity**

The distributions of the Gini index of reactivity are plotted for both control (navy) and salt treated (orange) transcripts and transcript regions in both shoot (A-D) and root (E-H). Changes in Gini index of reactivity between conditions of each transcript or region are shown by lines between the distributions. A full statistical summary of these changes can be found in Table S.3.b. The data used in these analyses can be found in Tables D.1-D.8.

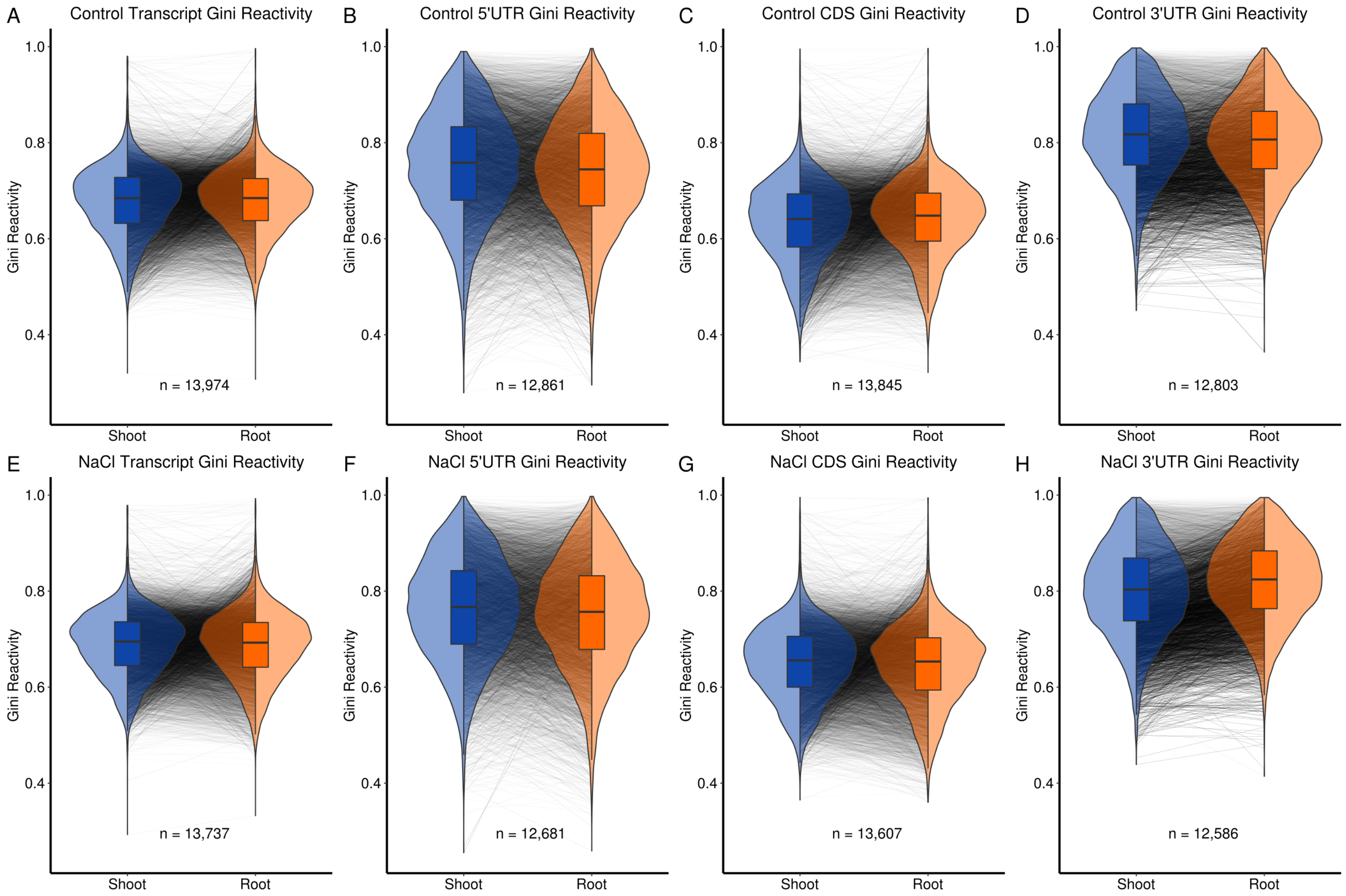


**Figure S.4 - The Effect of Salt Stress on Arabidopsis Inter-Tissue Gini Index of Transcript Reactivity**
The distributions of Gini index of reactivity are plotted for both shoot (navy) and root (orange) transcripts and transcript regions in both control (A-D) and salt (E-H) conditions. Changes in Gini index of reactivity between conditions of each transcript or region are shown by lines between the distributions. A full statistical summary of these changes can be found in Table S.3.b. The data used in these analyses can be found in Tables D.9-D.16.

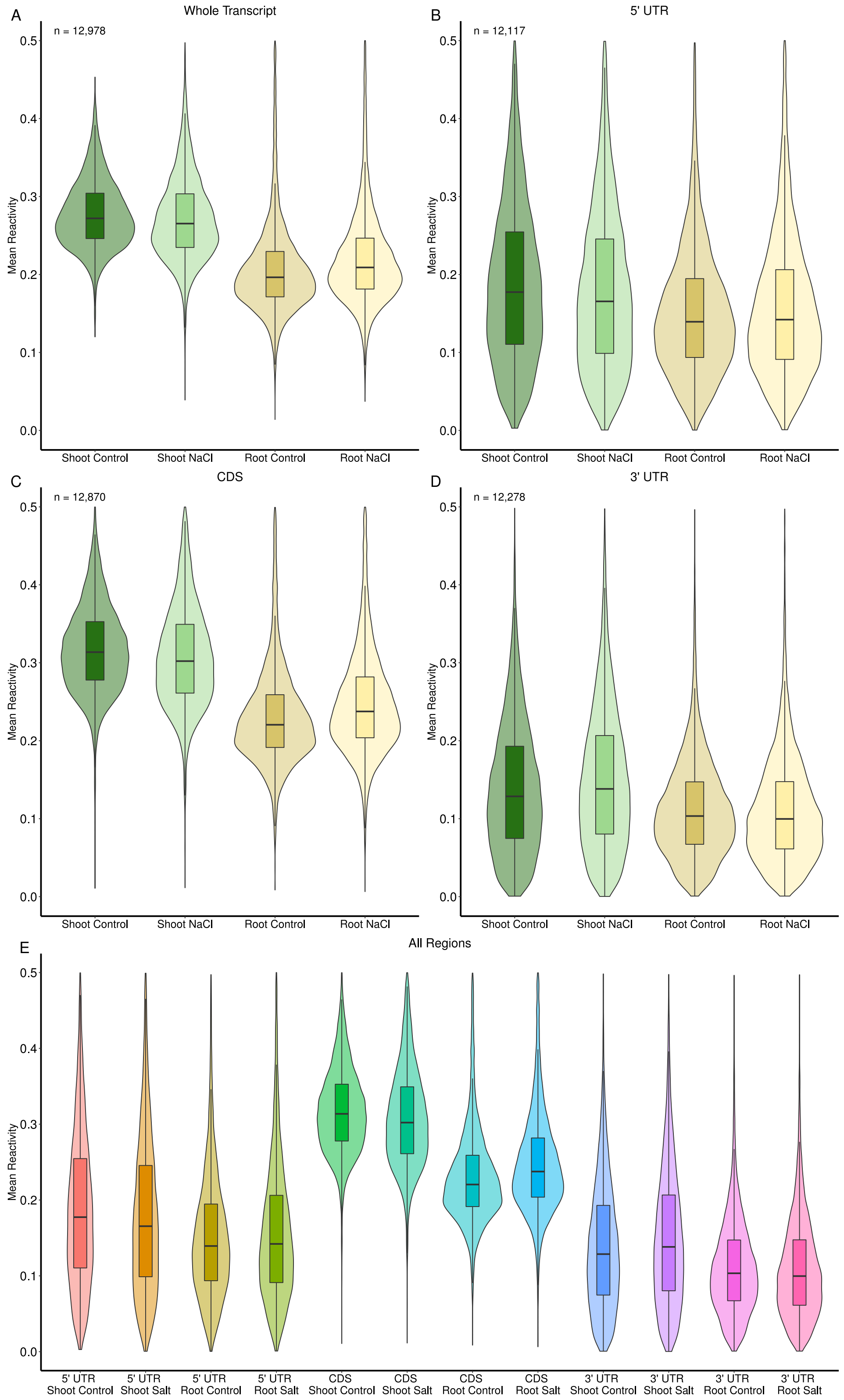


**Figure S.5 - The Effect of Salt Stress on Arabidopsis Transcript Mean Reactivity via ANOVA**

A-D) Two-way (mean~treatment*tissue) ANOVAs for transcript and transcript regions (Table S.5). E) Three-way (mean~treatment*tissue*region) ANOVA using region as a factor (Table S.5). Tissue and transcript region are more powerful determinants of mean reactivity than salt stress. The data used in these analyses are available in Tables D. 17-D.20.

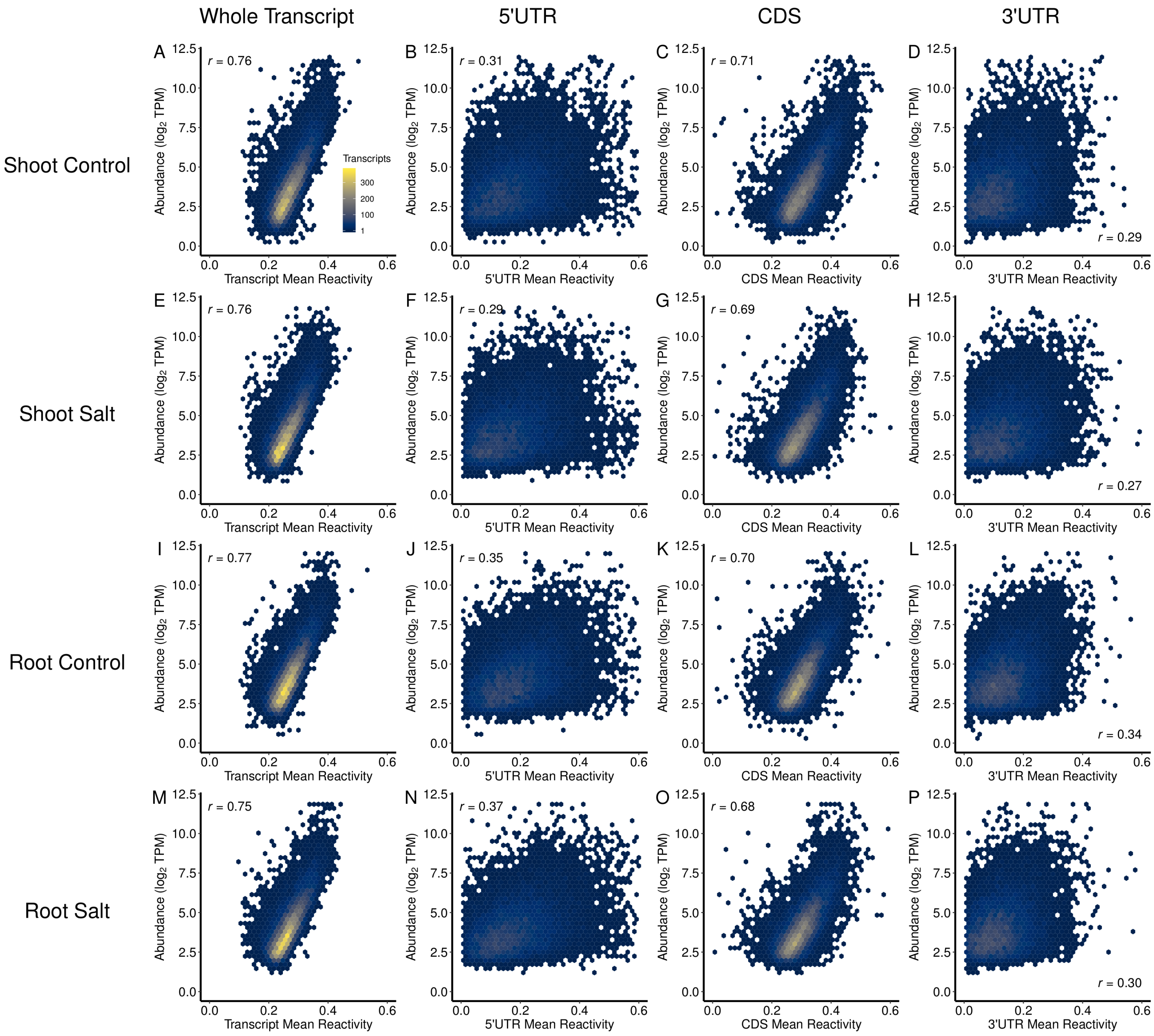


**Figure S.6 - Relationship between Reactivity and Abundance For Whole Transcript, 5’UTR, CDS, and 3’UTR with No Contrasts**

All statistics for the structurome of each tissue and condition were calculated independently, which allows the maximum number of transcripts to be resolved in each structurome as overlap of coverage between structuromes does not need to be considered. Additionally, this obviates the need for a common 2-8% scale, allowing for more precise quantification of trends. Panels for shoot control whole transcript (A) and root control whole transcript (I), mirror panels A and D from Figure 5 but with non-contrasted data, showing that the positive correlation between mean reactivity and abundance is present independently in each steady state structurome and transcriptome. Shoot control transcript regions (A-D), shoot salt transcript regions (E-H) root control transcript regions (I-L), root salt transcript regions (M-P). Recalculated reactivity data for whole transcript (R.1-R.4), 5’UTR(R.5-R.8), CDS (R.9-R.12), and 3’UTR (R.13-R.15) was used for all tests (Table S.6.c).

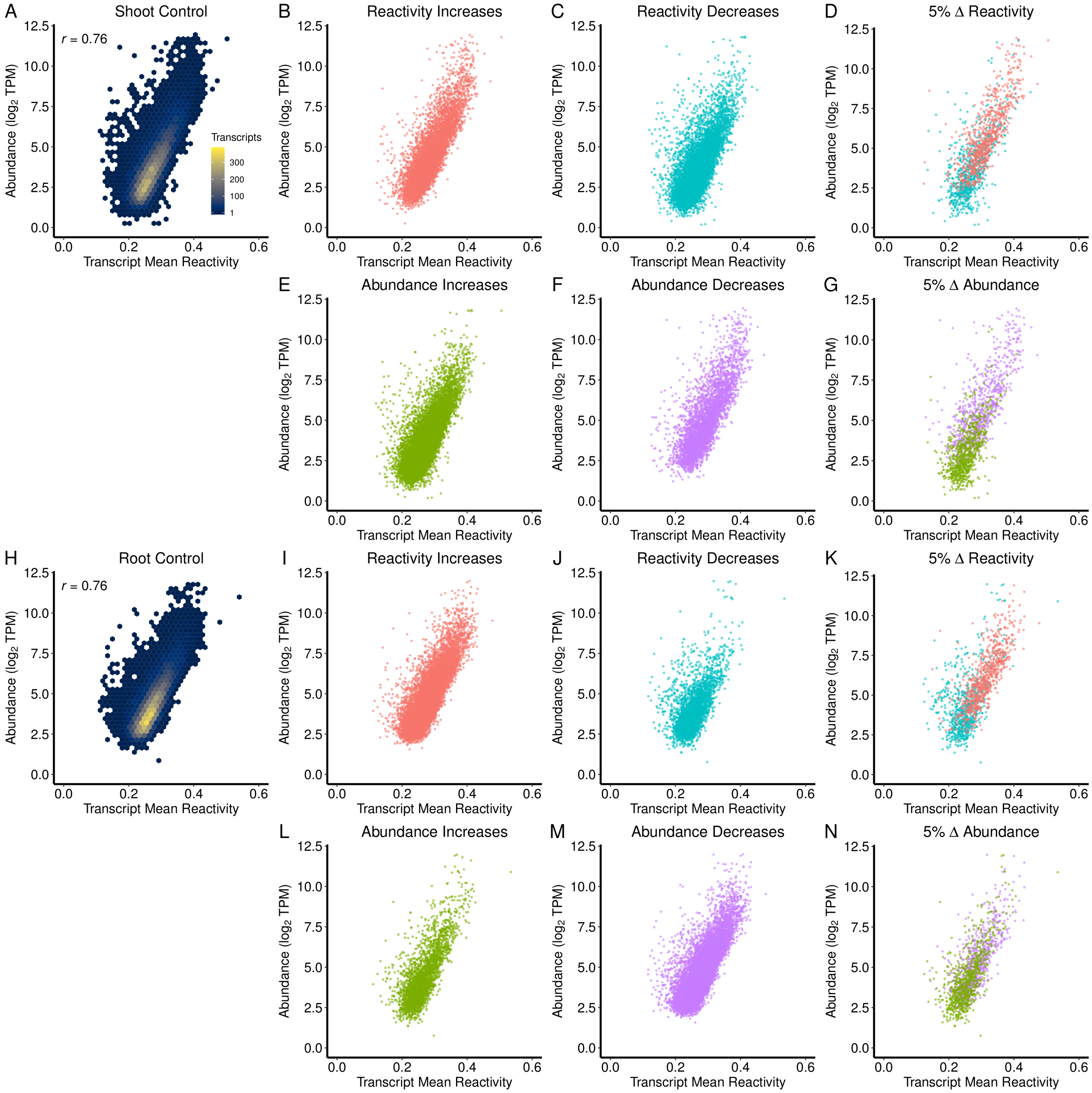


**Figure S.7 - Relationship between Reactivity and Abundance and its Modulation by Salt Stress, Extended**

Panels A, B, C of Figure 5 correspond to panels A, D, G of this extended figure; panels B and C show the full set of reactivity increases and decreases in shoot, and panels E and F show the full set of abundance increases and decreases in shoot. In the same fashion, panels D, E, F of Figure 5 correspond to panels H, K, N of this extended figure; panels I and J show the full set of reactivity increases and decreases in root, and panels L and M show the full set of abundance increases and decreases in root.

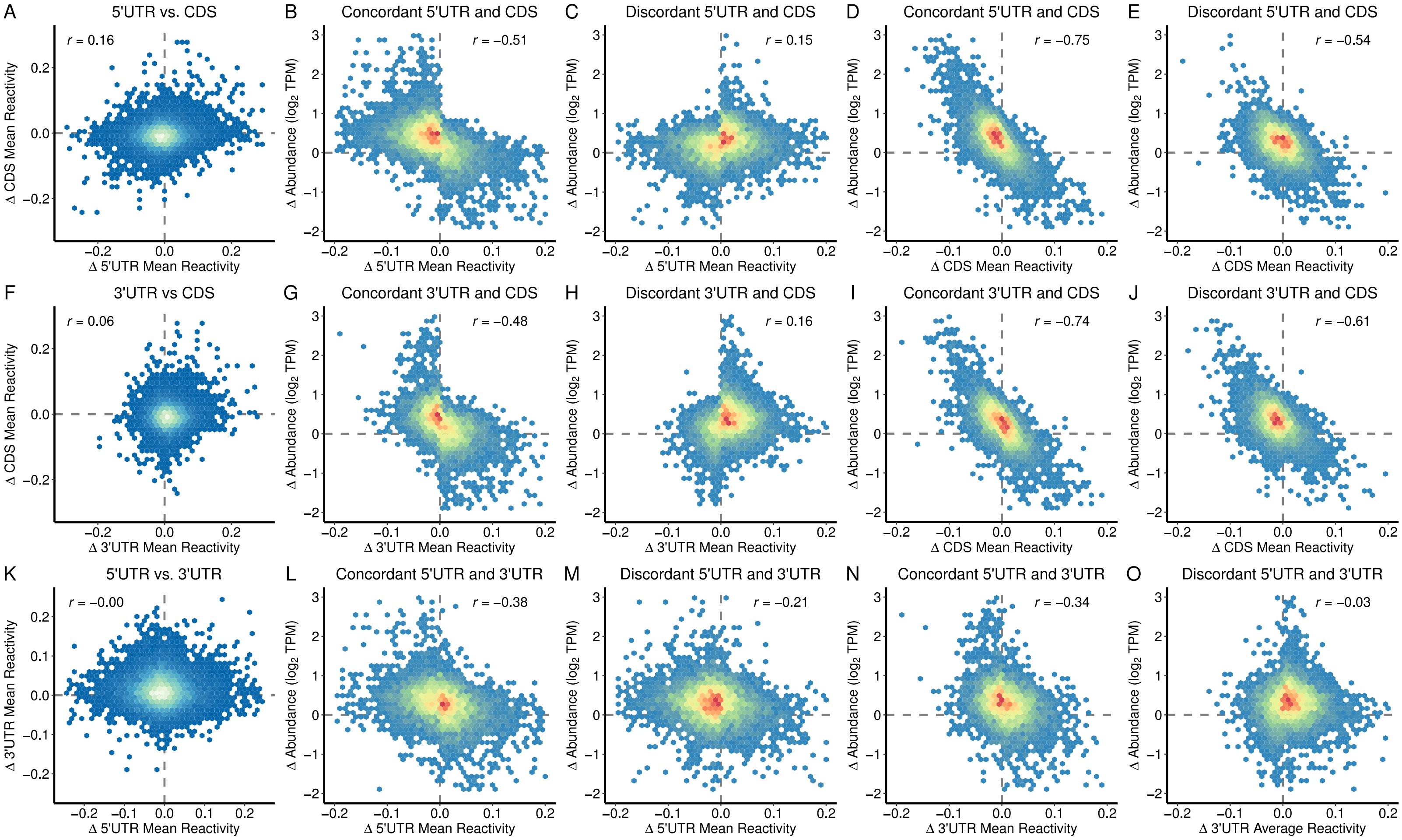


**Figure S.8 - Δ Reactivity Correlations, Concordant vs. Discordant, by Region, Shoot**

Column 1. For each two regions of each transcript resolved in our shoot contrast, we performed a Δ mean reactivity vs. Δ mean reactivity to test for a relationship between the regions' structural change; no such relationship exists (Table S.7.a). Columns 2 and 4. Transcript regions that concordantly change with another region have stronger Δ reactivity to Δ abundance correlations. Columns 3 and 5. Likewise, discordance tends to decrease the strength of the Δ reactivity to Δ abundance correlation. (Table S.7.b)

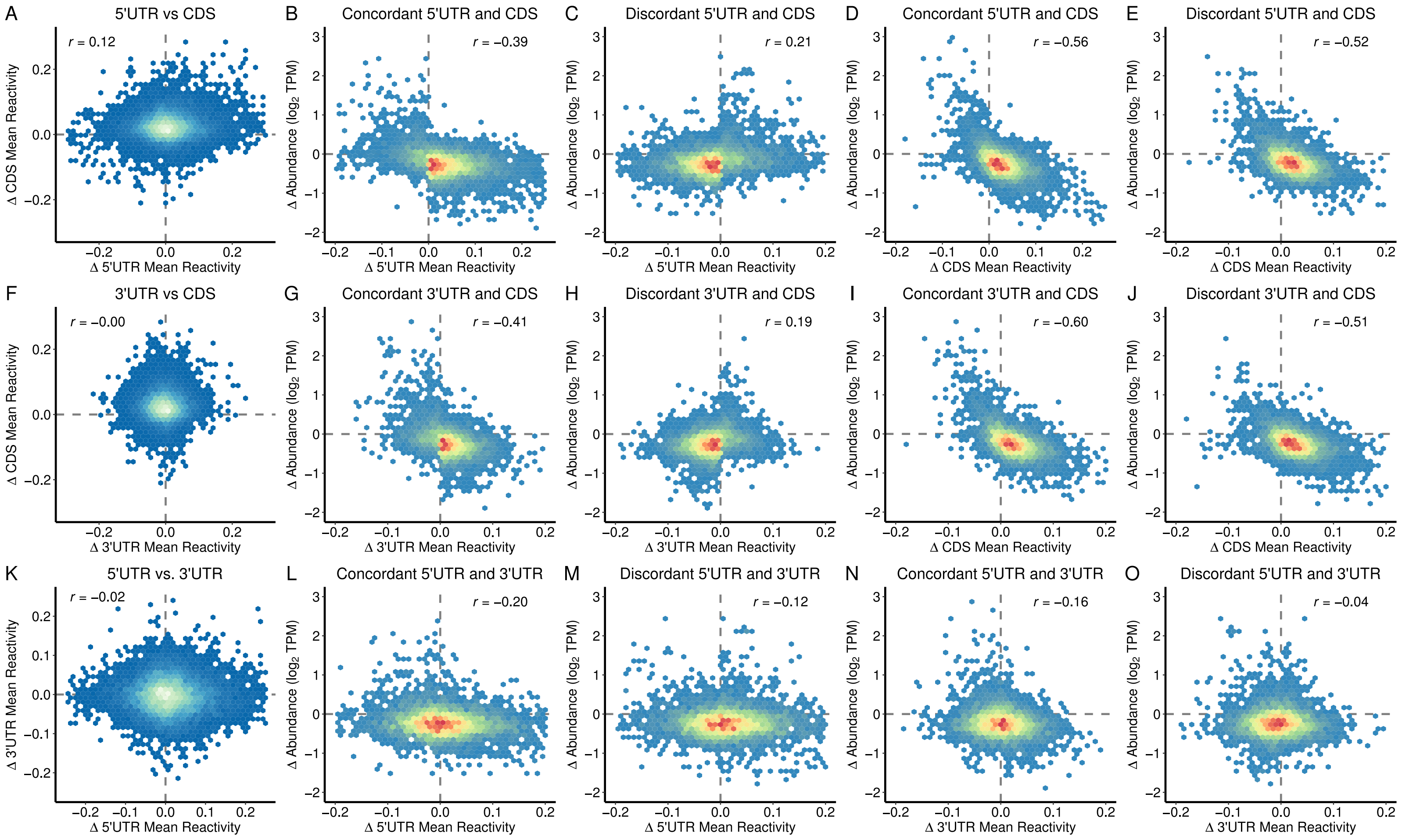


**Figure S.9 - Δ Reactivity Correlations, Concordant vs. Discordant, by Region, Root**

Column 1. For each two regions of each transcript resolved in our root contrast, we performed a Δ mean reactivity vs. Δ mean reactivity to test for a relationship between the regions' structural change; no such relationship exists (Table S.7.a). Columns 2 and 4. Transcript regions that concordantly change with another region have stronger Δ reactivity to Δ abundance correlations. Columns 3 and 5. Likewise discordance tends to decrease the strength of the Δ reactivity to Δ abundance correlation (Table S.7.b).

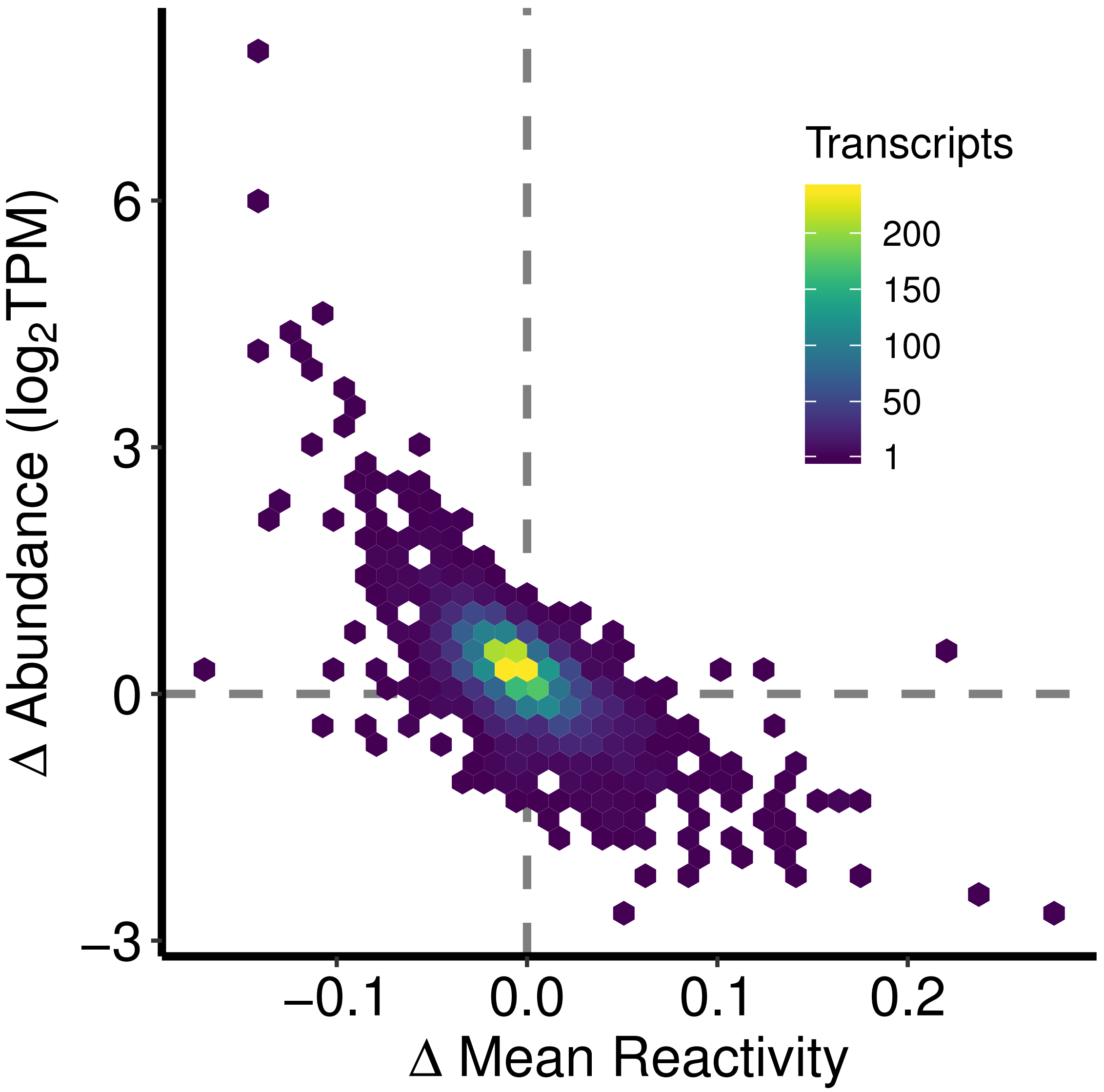


**Figure S.10 - Transcripts lacking m6A modification show the Inverse Relationship between Change in Reactivity and Change in Abundance**

All transcripts of genes that had one or more m6A peak in Anderson et al. 2018 were removed from our shoot data. The remaining data set was then plotted as in our Δ reactivity and Δ abundance plots, showing that the inverse correlation of Δ reactivity and Δ abundance is retained in transcripts with no detected m6A modifications.

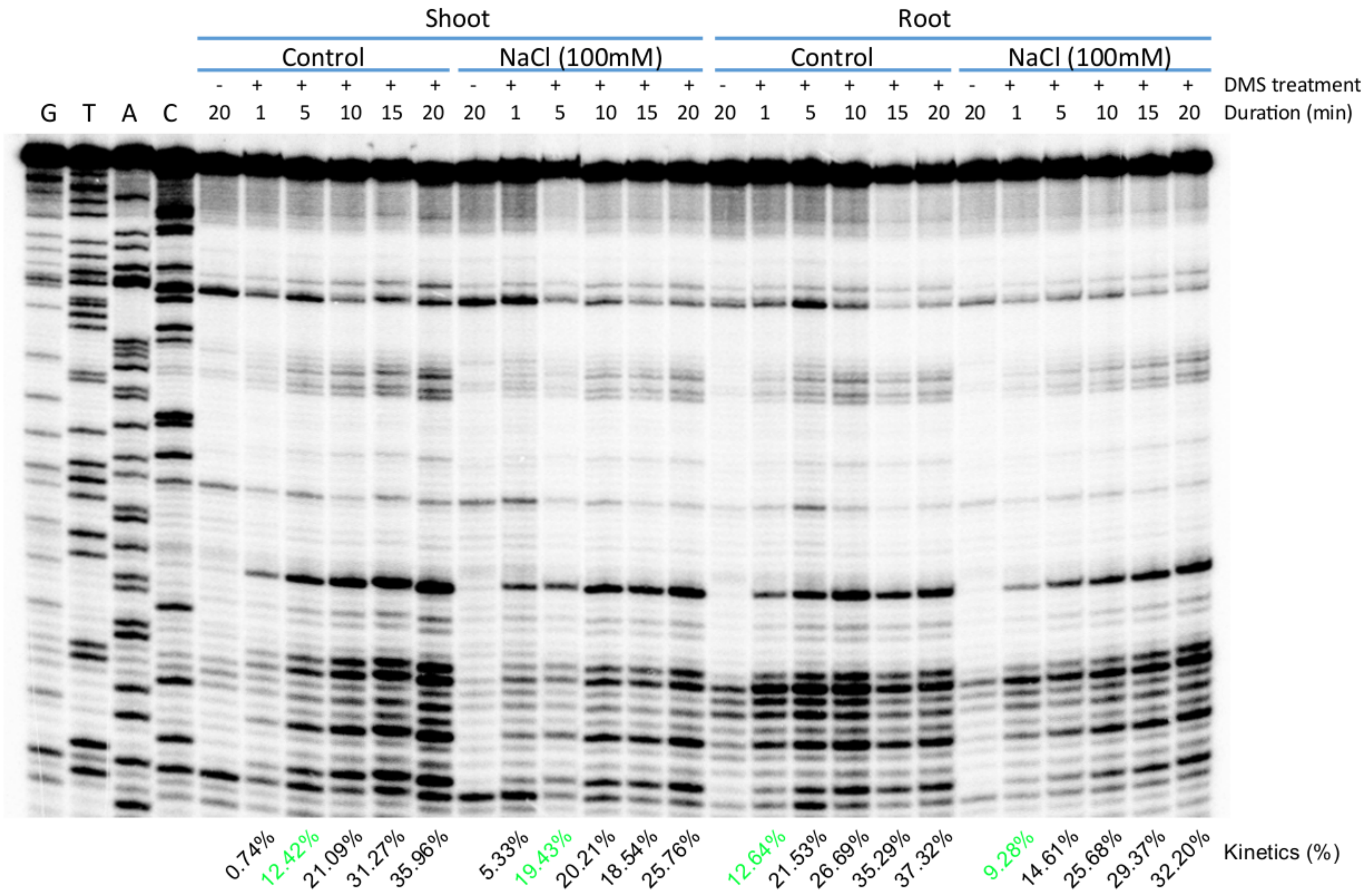


**Figure S.11 – Gel-based Assay for Single Hit Kinetics Determination**

Time course of in vivo DMS modification of Arabidopsis 18S rRNA in root and shoot tissue from control and salt-stressed plants. The extent of DMS modification was read out by gel-based probing. Single hit kinetics, defined as on average not more than one DMS modification per RNA molecule, was used as the criterion to determine the DMS treatment durations. A modification rate of 10-20% is consistent with the expected Poisson distribution of single-hit modification (Spitale et al. 2015). For each lane, modification rate was calculated as: [(the signal intensity of the full length band) / (the signal intensity of whole lane) x 100%]. Resultant optimal durations of DMS treatment are indicated in green font below the gel.
